# Expanding the *Bacteroides* synthetic biology toolkit to develop an *in vivo* intestinal malabsorption biosensor

**DOI:** 10.1101/2025.03.18.644028

**Authors:** Giselle McCallum, Juan C. Burckhardt, Jerry He, Alice Hong, Laurent Potvin-Trottier, Carolina Tropini

## Abstract

The human gut is a highly dynamic physical environment where perturbations—including factors such as acidification, oxygenation, and particle concentration (osmolality)—can influence microbiota composition and contribute to disease states. Understanding gut environmental changes is essential for advancing diagnostic and therapeutic strategies for gut health. However, non-invasive methods for continuous monitoring remain limited. The bacterial gut microbiota represents a powerful platform for continuous, non-invasive biosensing technologies for the gut environment, with genetically tractable commensal species like *Bacteroides thetaiotaomicron* (*B. theta*) emerging as promising hosts for engineered biosensors. However, the availability of genetic tools for precise, modular environmental sensing and reporting control in *B. theta* remains limited. Here, we present an expanded genetic engineering toolkit for *B. theta* that enables precise, fluorescence-based environmental sensing of the gut environment. This toolkit includes i) three libraries of orthogonally inducible promoters capable of driving fluorescence expression, ii) a DNA-based system to tune repressor activity in *B. theta*, iii) a resulting modular transcriptional reporter circuit that integrates native promoter activation with fluorescent outputs, and iv) characterization of a novel plasmid integration mode in *B. theta*. To demonstrate its utility, we engineered biosensors for gut malabsorption, a condition characterized by increased luminal osmolality. Using identified osmolality-responsive native promoters from *B. theta*, we made biosensors capable of detecting changes in gut physiology through graded fluorescent outputs. These biosensors were validated both *in vitro* and *in vivo* using a murine model of laxative-induced malabsorption, where they enabled continuous, long-term, non-invasive monitoring of single-cell response from fecal samples with sensitivity to subclinical malabsorption levels. By expanding the genetic toolkit for *Bacteroides* and demonstrating its use in a physiologically relevant context, this approach highlights the potential of engineered gut bacteria as a monitoring platform for diverse gut health applications. This work advances strategies for microbial biosensing and positions gut commensals as key players in next-generation diagnostic methods.

## Introduction

The human gut is a dynamic and complex ecological niche characterized by diverse physical environments^1^. These heterogeneous environments are the result of a combination of gradients in physical factors, such as pH, oxygen, and osmolality, that vary significantly throughout the gastrointestinal (GI) tract^2^. Aberrations in the physical environmental factors in the gut can contribute to, or result from, gastrointestinal diseases^2^. Understanding how changing physical parameters relate to disease states is critical for advancing diagnostic and therapeutic approaches to gut health. Currently, continuous, sensitive, and non-invasive monitoring of physical conditions in the gut is not possible due to technological limitations. Existing non-invasive methods, such as stool sample analyses (e.g., Bristol Stool Scoring, fecal calprotectin), are bulk measurements which are not sensitive to subclinical symptoms. Conversely, intestinal biopsies can be highly sensitive to local intestinal damage, but are limited to single time-point measurements and require invasive procedures such as endoscopies and colonoscopies with preparatory steps (i.e., bowel preparation) that alter the interrogated gut environment^1^. These challenges highlight the need for new technologies that can sensitively and continuously assess the gut environment non-invasively.

The bacterial microbiota offers a promising platform for detecting and reporting on the gut environment. Gut bacteria have evolved genetic mechanisms to detect and adjust to the dynamic changes encountered in their intestinal habitat. The transcriptional mechanisms employed by these microbes in response to environmental factors are promising options for developing microbiota-based strategies to monitor changes in gut conditions. Thanks to their small size and high sensitivity, bacteria can detect and respond to changes in environmental conditions along micron-scale gradients that are not detectable by most solid-state diagnostic tools. By engineering bacteria to produce reporter proteins (e.g., fluorescence, luminescence, or color signals) when physiological responses to an environmental cue are activated, biological sensors (biosensors) can be developed to report on the state of the gut environment non-invasively and potentially prior to the development of clinical symptoms.

In the context of the gut, *Bacteroides thetaiotaomicron* (*B. theta*) is a promising chassis for gut biosensors and engineered probiotics. Importantly, unlike *Escherichia coli*, *B. theta* is capable of long-term colonization at high abundance throughout most of the gut, including in intestinal crypts^3–5^. The development of *B. theta* synthetic biology tools has enabled precise gene expression tuning^6–9^, genetic memory systems^7^, user-controlled colonization and containment strategies^4,10^, and constitutive fluorescent protein expression^3^, advancing its engineering potential and application as a live probiotic in humans^10^.

However, a critical challenge remains: unlike *Escherichia coli*, native *B. theta* promoters cannot drive strong fluorescent protein expression^3,7,8^, limiting the possibility of single-cell interrogation of transcriptional activity *in situ* in this bacterium. Previous studies have relied on luminescence as a reporter of gene expression in *B. theta*^7–9^. However, luminescence is unable to be resolved at the single-cell level and is therefore limited to measuring population-average response. Luminescence reporters also cannot be multiplexed to measure multiple signals and are therefore more challenging to normalize. In contrast, single-cell analysis of transcriptional activity with fluorescence-based methods like flow cytometry or imaging can allow the identification of diverse responses within a population in the gut^11^. Using flow cytometry allows rapid screening of high numbers of cells and could therefore also enable detection of biosensor response from a population with limited colonization levels. Fluorescent tools for gut commensals like *B. theta* could therefore address the need for single-cell measurements of transcriptional activity in the gut.

Here, we addressed the need for sensitive, single-cell, fluorescence-based *B. theta* tools by developing a genetic toolkit for *B. theta* and using it to create a novel fluorescence-based transcriptional reporter system that enables expression reporting from native promoters. Starting from a previously discovered strong phage promoter capable of driving constitutive fluorescence expression in *B. theta*^3^, we engineered and characterized three libraries of repressible promoters that can drive tunable GFP expression with small-molecule inducers. We then identified native promoters upregulated under a variety of common stress conditions encountered in the gut with transcriptomics. Finally, we combined strong inducible promoters from our library with native inducible *B. theta* promoters, a TetR-based inverter module, and a DNA-based system to tune repressor activity^12^ to construct a modular biosensor circuit that reports native promoter expression via fluorescent outputs.

To demonstrate the utility of our inverted transcriptional reporter, we constructed and implemented biosensors to detect laxative-driven malabsorption that results in increased intestinal osmolality, using identified polyethylene glycol- and osmolality-responsive native *B. theta* promoters. In the gut, malabsorption is a clinical manifestation of common gastrointestinal pathologies, including inflammatory bowel disease (IBD)^13^, celiac disease^14,15^, small intestinal bacterial overgrowth (SIBO)^15^, food intolerances^15–17^, and enteric infections^15,18,19^. Our biosensors responded to both increasing osmolality levels *in vitro* and *in vivo* to laxative-induced malabsorption conditions in mouse models, as measured from single-cell fluorescence levels from fecal and cecal samples. Importantly, this technology enabled continuous detection of malabsorption severity from solid feces, even before overt clinical symptoms manifest. Our modular *B. theta* biosensors provide a platform for long-term, non-invasive, and continuous reporting of environmental conditions in the gut while significantly expanding the genetic engineering toolkit for this emerging model gut bacterium.

## Results

### Design rationale of a modular, inverted transcriptional reporter for *B. theta*

The inability of native *B. theta* promoters to drive high levels of fluorescent proteins limits the output of existing transcriptional reporters to luciferase or other enzymes^7^. As native promoters are capable of driving repressor expression to a functional level, we sought to build a circuit that links the output of a synthetic repressible promoter to the induction of a native promoter via a repressor-based inverter module (Figure 1A)^20,21^. Existing synthetic promoters have been engineered for *B. theta* by adding repressor binding sites to constitutive promoters, but these promoters cannot drive fluorophore expression^6–8^. Therefore, to achieve a repressible fluorescence output, we modified the previously discovered *Bacteroides*-targeting phage promoter P_BfP1E6_, which drives strong constitutive GFP expression in *B. theta*^3^. We engineered this promoter to be modulated by three different repressor proteins by adding one or two of the corresponding repressor operator sites. In this circuit design, native response to an environmental stimulus thus leads to a reciprocal increase in repressor expression, which decreases GFP expression levels in the cell (Figure 1A). To account for variations in overall cellular gene expression levels across growth phases, this circuit includes a constitutively expressed mScarlet-I3 red fluorescent protein^22^ (RFP) as a reference fluorophore. Native promoters can then be inserted modularly into the circuit using Golden Gate assembly^23^ to allow for easy construction of multiple biosensor circuits.

**Figure 1.**
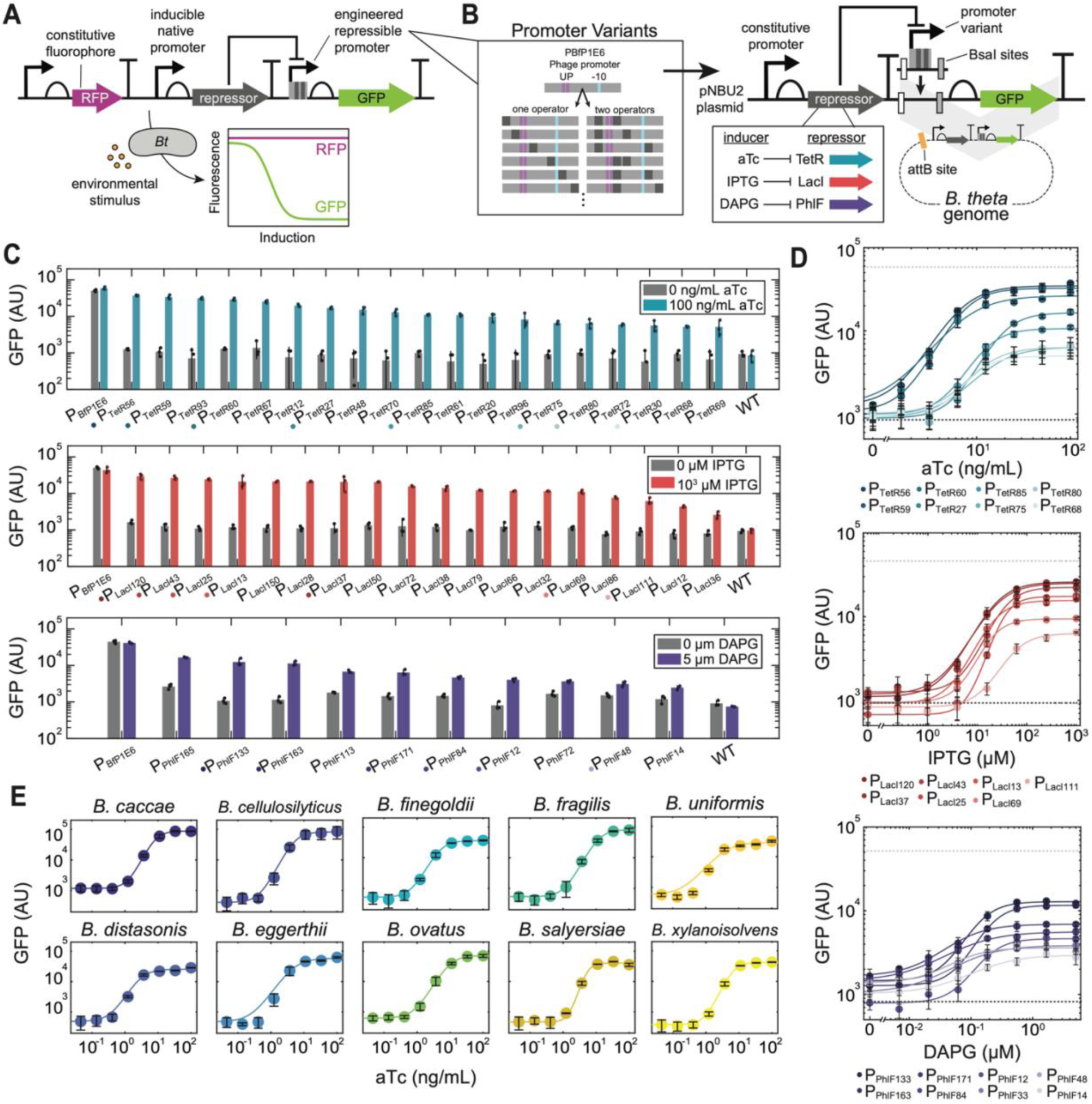
Engineered phage promoters enable repressible expression of GFP in *Bacteroides*. A) Design of an inverted, fluorescence-based transcriptional reporter circuit for *B. theta* (*Bt*), in which GFP expression is repressed by the induction of a native promoter expressing a repressor protein. B) Design and construction of small-molecule inducible promoter libraries in *B. theta*. C) Activity of promoter variants measured by GFP expression in uninduced (base media) and induced conditions (100 ng/mL aTc, 10^3^ μM IPTG, or 5 μM DAPG, respectively). D) Dose-response curves for 8 promoters from each library measured by GFP expression. Inducer concentrations were as follows: 2-fold serial dilutions starting at 100 ng/mL aTc; 4-fold serial dilutions starting at 10^3^ µM IPTG; and 3-fold serial dilutions starting at 5 μM DAPG. Dark and light dotted lines represent the WT and phage promoter (P_BfP1E6_) GFP levels under maximum induction levels, respectively. Fit lines represent a Hill function fit to dose-response curve data (See Supplementary Table 1 for fit parameters and goodness of fits). E) Dose-response curves for the P_TetR56_-GFP construct in 10 different *Bacteroides* species in 2.5-fold serial dilutions starting at 100 ng/mL aTc. Fit lines represent a Hill function fit to dose-response curve data. C-E: Error bars represent the standard deviation from n = 3 biological replicates (unique colonies) of each promoter. AU: Arbitrary units

Importantly, an inverted transcriptional reporter offers several advantages to direct reporters for biosensor applications. As the reporter is constitutively expressed in the absence of a stimulus, the strong baseline signal ensures that even small changes in repression can be detected, improving sensitivity. Additionally, in the case that native promoter induction is weak, direct reporters can suffer from limited signal amplification, resulting in non-detectable reporter expression. In contrast, inverted systems can use repressor proteins to translate native gene expression to amplified, non-linear decreases in the expression of a reporter, allowing a larger detectable dynamic range and clearer distinctions between active and inactive states.

### A library of small-molecule inducible promoters facilitates tuneable GFP expression levels in *B. theta*

As our biosensor design relies on controlling GFP expression indirectly through the expression of a cognate repressor protein (Figure 1A), we sought to engineer a promoter that i) is tightly regulated by a repressor protein not naturally present in *B. theta*, and ii) has a large dynamic range of expression. To achieve this, we constructed three libraries of promoters regulated by TetR, LacI, or PhlF repressors (consisting of 144, 245, and 131 promoters, respectively) by inserting one or two copies of the corresponding operator sites throughout the P_BfP1E6_ promoter in locations that did not disrupt the UP elements or −7/10 consensus sequence^3^ (Figure 1B). Promoters were inserted upstream of a super folder GFP gene in a Golden Gate-adapted pNBU2 *Bacteroides* conjugation plasmid^3,24^. Constructs were conjugated into *B. theta* and integration into the genome at one of two known integration sites at the 3’ end of the serine tRNA genes (attBT2-1 or attBT2-2)^24^ was verified with PCR (Methods). To identify promoters with a large dynamic range and low baseline expression, we measured GFP expression from 192 individual colonies from each library by growing them in the presence or absence of the appropriate small molecule inducer (TetR: anhydrotetracycline [aTc]^8^, LacI: ß-D-1-thiogalactopyranoside [IPTG]^7^, or PhlF: 2,4-diacetylphloroglucinol [DAPG]^25^). Promoter variants in all libraries displayed a wide range of baseline and maximal expression levels, indicating a differing effect of operator site position on repression efficiency and maximum promoter strength (Supplementary Figure 1A). The parent P_BfP1E6_ promoter did not respond substantially to any inducer, while each library contained multiple promoter variants of interest which showed strong repression and a large dynamic range (i.e., low GFP expression at baseline, and strong induced GFP expression)(Supplementary Figure 1A). Up to 19 unique functional variants from each library with a range of induction levels were further characterized (Figure 1C) and sequenced (Supplementary Table 1). Based on the two criteria defined above, the best promoter candidates from each library were P_TetR56_, P_LacI43_, and P_PhlF133_, which showed strong repression that was close to the autofluorescence of wild-type cells (2.4%, 2.9%, and 2.1% of the parent promoter, respectively, compared to 1.6% for wild-type) (Figure 1C). These promoters also displayed a large dynamic range (29.4-, 20.8-, and 11.6-fold change expression, respectively) and strong maximal expression (63.5%, 61.2%, and 40.1% of the parent P_BfP1E6_ promoter, respectively). These expression levels are substantially higher than strong native promoters, including P*_rpoD_* and P_rRNA_, which only showed up to ∼20% of the P_BfP1E6_ promoter (Supplementary Figure 1B). Promoters from our libraries also showed stronger induced expression levels than previously engineered TetR- and LacI-regulated repressible promoters for *B. theta* (P1T_DP_^8^ and P ^7^), which only showed 2.7- and 1.5-fold activation, respectively (Supplementary Figure 1C). To further characterize highly repressed promoters from each library, we performed dose-response curves for promoter variants by growing them in a range of concentrations of their respective small molecule inducers. Dose-response curves from promoters in all libraries were fit well with a Hill function typical of small molecule inducible promoters (r^2^ > 0.9; Supplementary Table 1) and displayed a range of values for maximum expression levels (*V_max_*), induction thresholds (*K_m_*), and Hill coefficients (*h*) (Figure 1D; Supplementary Table 1).

Given the robust performance of the P_TetR56_, P_LacI43_, and P_PhlF133_ inducible promoters, we tested for orthogonality between the induction systems, as this is a critical feature for the construction of complex synthetic circuits^7^. We measured the degree of cross-induction between strong promoters in our libraries by growing several promoter variants from each library in aTc, IPTG, and DAPG, and compared GFP levels to uninduced controls. Promoters were only induced in their cognate inducer (Supplementary Figure 1D), indicating that our induction systems are highly orthogonal and could be implemented in complex circuits with minimal crosstalk.

The portability of these tools to other *Bacteroides* would support genetic engineering in other important gut commensals of interest. We therefore tested whether our promoters were functional in an expanded host range. We conjugated ten *Bacteroides* species with the P_TetR56_-GFP plasmid and measured GFP expression in response to aTc induction. Induction followed the expected Hill dynamics and showed a strong increase in expression levels in all species (Figure 1E). These results suggest that P_TetR56_, along with other promoters in our libraries, can support strong, tunable gene expression across diverse *Bacteroides*, highlighting its potential as a broadly useful genetic part.

Our engineered promoter libraries expand the number of inducible promoter variants available for *B. theta* and offer a range of induction dynamics for the construction of complex synthetic circuits. To the best of our knowledge, these represent the first inducible promoters that can drive fluorescent protein expression in *Bacteroides*, opening the possibility for synthetic circuits reported by fluorescence in these gut bacteria. Given its high *V_max_* and tight baseline repression, we selected P_TetR56_ and TetR as the repressible promoter and cognate repressor protein to be implemented in our inverted *B. theta* biosensor circuit.

### Gene expression analysis of *B. theta* in physiological stress conditions identifies upregulated promoters that are perturbation-specific

Having engineered a tunable promoter with a high dynamic range to form the backbone of an inducible reporter system, we next sought to identify native *B. theta* genes that are regulated uniquely and robustly in response to changing environmental conditions. Promoters from these genes could be used to drive TetR expression in our circuit, such that their induction leads to lower expression of GFP from the P_TetR56_ promoter. To this end, we performed transcriptomic profiling of *B. theta* grown *in vitro* under various stress conditions that occur in the gut (namely, pH, oxidative, temperature, and osmotic stress). After identifying promoters that respond specifically to each of these stressors, we then focused our analysis on osmotic stress as a key disrupted factor in multiple intestinal diseases that could be detected with *B. theta* biosensors.

We first performed RNA-seq on *B. theta* cultures grown in complex base medium (∼320 mOsm/kg, pH 7, 37 °C), or in stress conditions (pH 5.8, 42 °C, or 100 µM H_2_O_2_). For osmotic stress, we tested response to four osmolytes (sodium chloride [NaCl], sorbitol, lactulose, and polyethylene glycol [PEG]) at two osmolalities representative of intestinal levels during malabsorption (∼700 and ∼900 mOsm/kg)^26^. These compounds span common types of osmolytes that can be found in the gut from dietary or pharmaceutical sources (i.e., salts, sugar and sugar alcohols, and commercial laxatives, respectively). Overall, most conditions cluster away from the baseline samples in Principal Component Analysis of RNA-seq data (Supplementary Figure 2A), indicating a shift in the gene expression profile of *B. theta* in response to these perturbations. Within osmotic stress conditions, each osmolyte resulted in a unique transcriptome signature (Supplementary Figure 2A), supporting the use of multiple osmolytes to identify genes that are broadly responsive to osmolality.

To identify genes that were significantly upregulated in each stress condition, we performed differential expression analysis for each stress condition compared to base media (Supplementary Table 2). All conditions tested led to significantly increased expression of at least some genes (Figure 2A), and each stress condition showed a set of uniquely upregulated genes (Figure 2B). These uniquely responsive genes may be promising for use in biosensors targeted to detecting specific stress conditions in the gut. We also identified 247 genes that were upregulated in all the conditions tested, which could be of interest in the context of a broad stress sensor. We focused our search on osmolality-responsive genes to develop an intestinal malabsorption biosensor. Specifically, we looked for genes that were not induced under other stress conditions tested and were activated proportionally to increased concentration of osmolytes. We identified 155 genes that were significantly upregulated in at least one osmolyte at ∼900 mOsm/kg (around the highest level of malabsorption measured in the mouse gut)^26,27^, but no other stress condition (Figure 2B). Using the reference *B. theta* genome (GCF_000011065.1) to annotate differentially regulated genes, we found the 10 most strongly upregulated genes in at least one osmotic stress condition to be broadly involved in biosynthetic pathways, signal regulation, and transmembrane protein transport (Figure 2C,D; Supplementary Table 3). Many of these genes were also upregulated at ∼700 mOsm/kg (representative of moderate malabsorption in the gut) and showed a graded induction in response to increasing osmolality levels in more than one osmolyte, which is a desirable feature for building a broad biosensor for osmotic stress (Figure 2D). Our transcriptomics analyses provide a valuable map of transcriptome response to osmolality and other common stressors in the gut. Altogether, our analyses identified native *B. theta* promoters that respond strongly and specifically to a range of environmental perturbations, providing a useful resource to select gene candidates for the construction of biosensors for environmental parameters in the gut.

**Figure 2.**
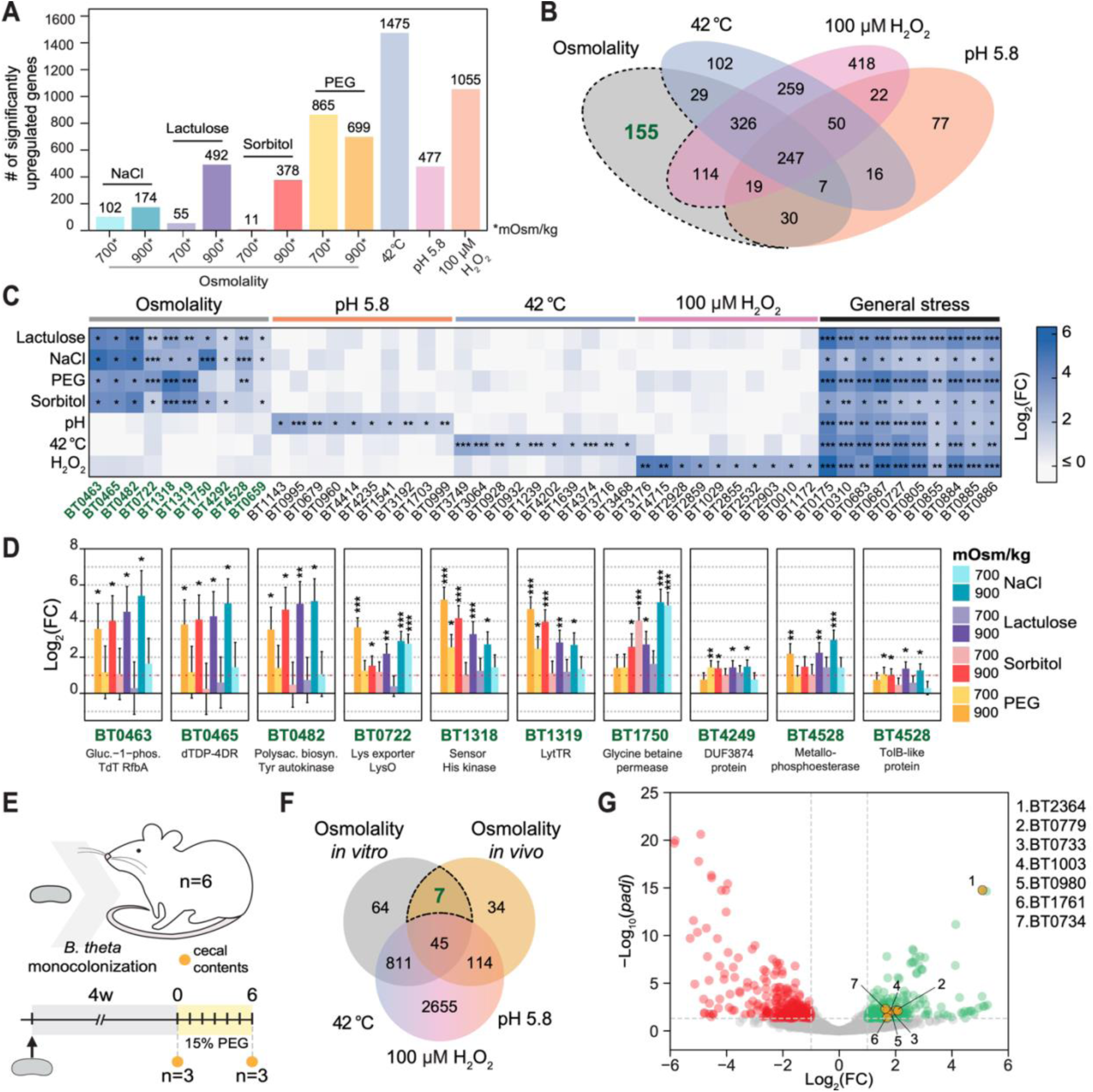
**Native *B. theta* promoters respond specifically to physical perturbations *in vitro* and to osmotic stress *in vivo*.** A) Significantly upregulated genes compared to base media (*padj* < 0.05 and Log_2_(FC)> 1) for every *in vitro* condition. B) Genes significantly upregulated compared to base media that are unique to specific stresses or shared between conditions (responsive to multiple stressors). Dotted line specifies genes unique to osmotic stress that were significantly upregulated in at least one osmolyte at high osmolality (∼900 mOsm/kg) C) Log_2_(FC) of the 10 most upregulated genes unique to each stress condition (osmolality, pH 5, 42 °C and 100 µM H_2_O_2_) and top 10 most upregulated genes present in all conditions (general stress). D) Log_2_(FC) of the top 10 genes from osmotic stress conditions at medium (∼700 mOsm/kg) and high (∼900 mOsm/kg) for all osmolytes tested. E) Experimental schematic for *in vivo* RNA-seq experiment. F) Genes significantly upregulated (*padj* < 0.05 and Log_2_(FC) > 1) compared to base media or untreated mice for *in vitro* and *in vivo* experiments, respectively, that are unique or shared between conditions tested. Dotted line highlights genes upregulated in at least one of the high osmolality (900 mOsm/kg) conditions *in vitro*, upregulated *in vivo*, but not upregulated in the general stress conditions. G) Volcano plot showing gene expression changes of all *B. theta* genes from the *in vivo* PEG experiment. Highlighted points are the 7 genes identified in Figure 2F. D: Error bars represent the standard error from *n* = 3 biological replicates. C,D: Statistics calculated in DESeq2^30^ pipeline (* *padj* < 0.05; ** *padj* < 0.01; *** *padj* < 0.001)

As a biosensor ultimately needs to respond in the gut environment, we next sought to identify additional genes that may be responsive to osmotic stress *in vivo*. PEG is a common laxative used to induce malabsorption and has previously been established as a malabsorption and osmotic diarrhea model in mice^26–29^. To characterize the *B. theta* transcriptional response to increased osmolality *in vivo*, we therefore mono-colonized germ-free mice with *B. theta* and treated them with PEG in drinking water for 6 days as previously shown (Figure 2E)^26^. We then measured intestinal osmolality as well as bacterial abundance and performed RNA-seq on the cecal contents of PEG-treated and control mice. PEG treatment resulted in an increased cecal osmolality to ∼800 mOsm/kg compared to the 400 mOsm/kg of control mice (Supplementary Figure 2B) and a 10-fold decrease in bacterial abundance (Supplementary Figure 2C). Differential expression analysis from RNA-seq showed that PEG treatment significantly shifted the overall *B. theta* transcriptome (Supplementary Figure 2D; Supplementary Table 4). To find promoter candidates to use in an osmolality biosensor, we identified 7 genes from this dataset that were i) upregulated during PEG treatment *in vivo*, ii) upregulated in at least one 900 mOsm/kg osmotic stress condition *in vitro,* and iii) were not upregulated in other stress conditions *in vitro*. (Figure 2F; Supplementary Table 3). These genes included outer membrane proteins, hypothetical proteins, a histidine kinase, glycoside hydrolases, and a polysaccharide lyase. Most of these genes showed 2- to 4-fold higher expression during PEG treatment (Figure 2G) except for BT2364, a TonB-dependent transporter that showed a ∼32-fold increase during PEG treatment (Figure 2G). Many of these genes were upregulated in PEG both *in vitro* and *in vivo* and some were significantly upregulated during growth in high osmolality induced by other osmolytes (Supplementary Figure 2E). The promoters identified *in vitro* and *in vivo* displayed specific activation in response to osmotic stress and graded induction across a range of osmolality, making them promising candidates for the construction of biosensors that respond precisely to changes in intestinal osmolality.

### Biosensors incorporating osmolality-specific promoters respond to osmotic stress *in vitro*

Having identified a selection of osmolality-specific *B. theta* genes, we next aimed to construct candidate osmolality biosensor circuits using promoters from these genes in our inverted transcriptional reporter design (Figure 1A). To screen for functional biosensors, we inserted the best 5 promoter candidates from genes identified in both our *in vitro* and *in vivo* RNA-seq analyses into our circuit upstream of the *tetR* gene, with the engineered P_TetR56_ promoter driving GFP (Figure 3A). As we ultimately required a fluorescent output that could be distinguished in fecal material, we changed the GFP in our reporter circuit to mGreenLantern (mGL), which is known to be brighter than sfGFP^31,32^. We then grew biosensor strains in base media (∼320 mOsm/Kg) and media adjusted to high osmolality (∼850 mOsm/kg) with PEG and measured cellular GFP and RFP levels. We calculated reporter activation as the fold decrease in the ratio of GFP to the constitutive RFP control (FC(GFP:RFP); activation) between base media and osmotic stress conditions. To verify that GFP levels are not decreased by osmotic conditions in the absence of promoter induction, we constructed a control strain in which no promoter is inserted upstream of TetR, and GFP is therefore not repressed (no promoter control [NP]).

**Figure 3.**
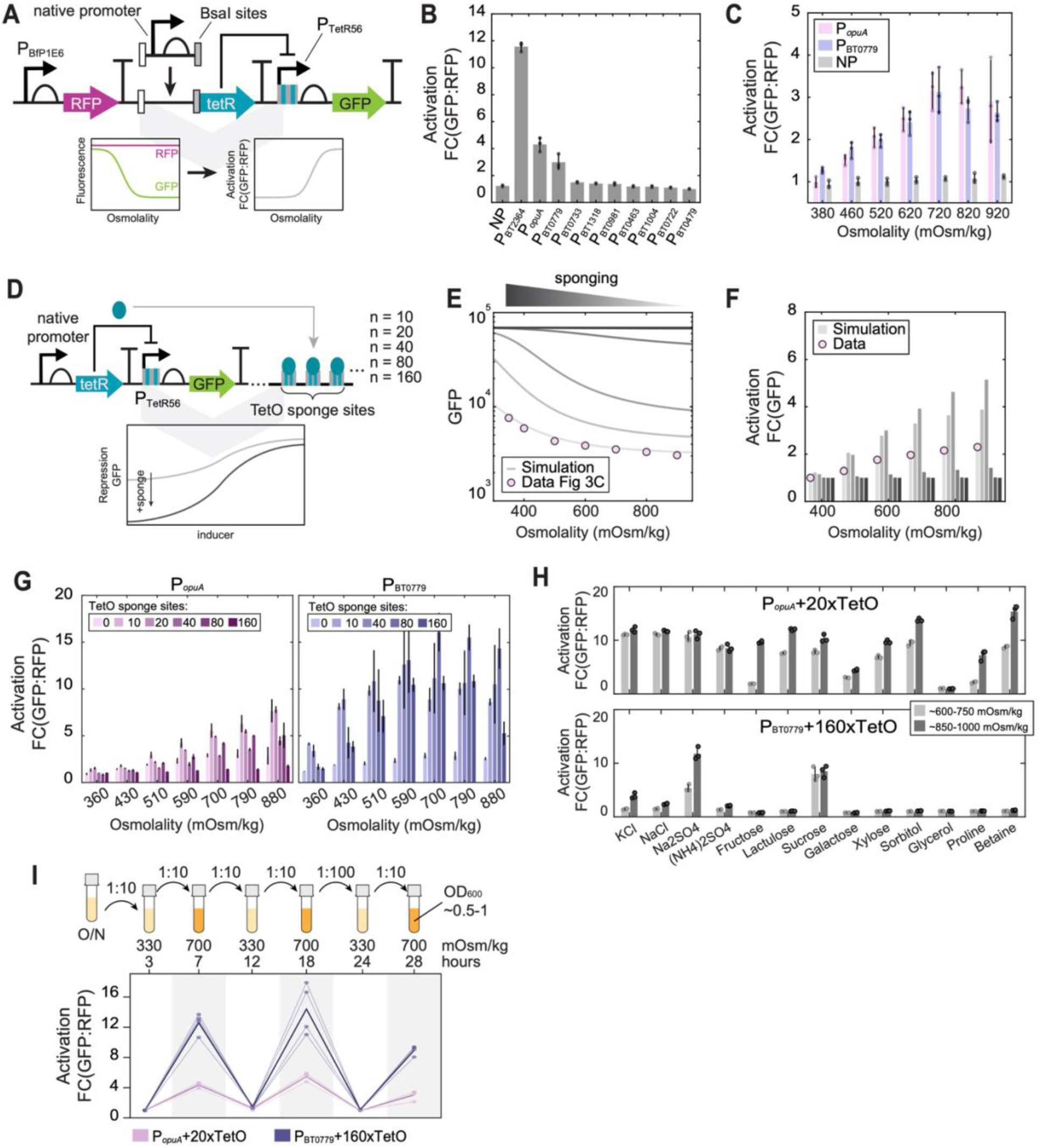
***In vitro* biosensor response is enhanced by TetR sequestration, responsive to multiple osmolytes, and reversible.** A) A modular Golden Gate entry vector facilitates the assembly of *B. theta* biosensors which can report on the activity of arbitrary native promoters. GFP is downregulated as the native promoter is induced, and a constitutive RFP expressed from P_BfP1E6_ allows for normalization. Biosensor activity is measured as the fold decrease in the ratio GFP to RFP (FC(GFP:RFP); activation) compared to baseline conditions. B) Induction response of a selection of candidate biosensors using different native promoters or no promoter (NP) grown in baseline media (∼320 mOsm/kg) or media adjusted to ∼850 mOsm/kg with PEG. C) Dose-response curves for the P*_opuA_* and P_BT0779_ biosensors in media adjusted to osmolality with PEG. D) Design of a biosensor with TetR ‘sponging’, which is predicted to lower baseline repression of GFP. E) Steady-state GFP levels from simulations of the inverted P*_opuA_*biosensor circuit with increasing degrees of TetR sponging. Model parameters were fitted to the PEG dose-response curve for the no-sponge P*_opuA_* biosensor shown in C (pink circles). F) Fold decrease in steady state GFP levels from simulations shown in E. G) Biosensor response to osmolality with genetically encoded TetR sequestration. P*_opuA_* and P_BT0779_ biosensors with 0, 10, 20, 40, 80, or 160 TetO binding sites present in the circuit were grown in media adjusted to specified osmolalities with PEG. H) Optimized P*_opuA_*+20xTetO and P_BT0779_+160xTetO biosensor response measured in media adjusted to moderate (∼600-750 mOsm/kg) or high (∼850-1000 mOsm/kg) osmolality with a variety of osmolytes. I) Activity of P*_opuA_*+20xTetO and P_BT0779_+160xTetO biosensors during three cycles of alternating growth in low (330 mOsm/kg) and high (700 mOsm/kg) osmolality media modified with PEG. B,C,G,H: Error bars represent the standard deviation from *n* = 3 biological replicates (unique colonies) of each biosensor. Fold decrease in GFP:RFP (activation) is calculated relative to those in baseline conditions (∼320 mOsm/kg).

Three biosensor candidates incorporating the promoters from the BT2364, *opuA* (BT1749-51), and BT0779 genes showed activation in response to increased osmolality *in vitro* (Figure 3B), while the NP control remained inactive (Figure 3B; Supplementary Figure 3A). The *opuA* operon is known to be involved in osmotic stress response in many other bacteria^33–35^. Conversely, BT0779 is a hypothetical protein that has homology to a *Bacteroides* HdeA/HdeB chaperone family protein. The P*_opuA_* and P_BT0779_ biosensors showed a 4.3- and 3.0-fold activation in media adjusted with PEG, respectively (Figure 3B). As expected, this change in GFP:RFP ratio was due to increased repression of GFP with increased osmolality in both biosensors, while RFP levels remained stable across conditions (Supplementary Figure 3A). In contrast, both the NP control and the remaining biosensors, except for P_BT2364_, showed minimal responses (<1.2-fold activation; Figure 3B; Supplementary Figure 3A).

BT2364 encodes a SusC/D homologue, suggesting its expression may be linked to carbohydrate utilization in the gut. When tested *in vitro*, the P_BT2364_ biosensor showed little activation in response to growth in various carbohydrates, except for sucrose (Supplementary Figure 3B). However, given that previous studies have found this promoter to be regulated by dietary carbohydrates^36,37^ and species-specific interactions in the gut^38^, we deemed it less suitable as an osmolality-specific biosensor and therefore focused on the other two candidate promoters. Interestingly, promoters upstream of putative LytS/LytTR-type histidine kinase– response regulator genes (P_BT0733_ and P_BT1318_) did not display an inducible response in our assays, despite the established role of these systems in mediating adaptation to osmotic and acid stress via signal transduction cascades^39^.

Taking the P*_opuA_* and P_BT0779_ biosensors as our most promising candidates, we aimed to further characterize the dose-responses and dynamic range of these reporters. To this end, we grew these strains in media ranging from ∼320 to ∼900 mOsm/kg, adjusted with PEG, and measured reporter activation. Both biosensors showed repression of GFP leading to graded biosensor activation in response to increasing osmolality, which reaches maximal activation at ∼700 mOsm/kg (Figure 3C; Supplementary Figure 3C).

The graded activation of the two biosensors to increasing levels of PEG-mediated osmolality *in vitro* suggested that their activity could also detect PEG-induced malabsorption in the gut. Before proceeding to *in vivo* studies, however, we sought to improve their dynamic range and tunability by addressing a key limitation observed *in vitro*.

### TetR sequestration improves biosensor performance *in vitro*

While both the P*_opuA_* and P_BT0779_ biosensors responded to increasing osmolality, we noticed that GFP levels were substantially lower than the NP control in baseline media (18.9% and 9.1% of the NP control, respectively) (Supplementary Figure 3C), suggesting that non-zero expression of TetR from the native promoters led to repression of GFP. Since our biosensor response depends on repression of GFP, we reasoned that low baseline GFP levels constrained the dynamic range of our sensors. We therefore hypothesized that sequestering excess TetR from our circuit would increase baseline GFP levels and improve the potential dynamic range of our reporters (Figure 3D). Indeed, previous synthetic circuits incorporating DNA-based repressor-binding “sponges” have been shown to decrease baseline levels of repressor^40^ and increase fold change of circuit outputs^12^.

To predict the effects of repressor sequestration on the dynamic range of our inverted transcriptional reporter circuit, we simulated our circuit using an ODE-based mathematical model (Methods). The model includes transcription and translation of TetR and GFP, and a TetR ‘sponging’ term that follows Hill ligand binding dynamics. After fitting model parameters (Supplementary Table 5) using dose-response curves for the P_TetR56_ promoter (Figure 1D) and the P*_opuA_* biosensor (Supplementary Figure 3C), we calculated steady-state GFP levels over a range of induction with varying degrees of sponging (Figure 3E). Including TetR sponging in our model improved the dynamic range of the system at all induction levels (Figure 3F), though over-sponging resulting in a lack of GFP repression at high induction levels was possible (Figure 3F).

To implement repressor sponging experimentally, we used both chemical and DNA-based sponging methods to sequester TetR in two experiments. We first sequestered TetR chemically using aTc to prevent binding to the TetO operator site, as this method offered a rapid method to evaluate the effect of TetR sequestration without the need for additional cloning. Here, we performed PEG-osmolality dose-response curves as above, but with a range of aTc concentrations supplemented into growth media (Supplementary Figure 3D). In agreement with modeling predictions, TetR sequestration using aTc improved the dynamic range of the biosensors. Specifically, both P*_opuA_* and P_BT0779_ biosensors reached a maximal activation (4.9- and 7.8-fold, respectively) at the highest osmolality tested (∼850 mOsm/kg) with 0.25 ng/mL aTc supplemented into the media (Supplementary Figure 3D). While these results confirm that TetR sequestration improved biosensor performance, a sponging system that does not require small molecule supplementation is preferable for *in vivo* biosensor implementation. Next, we therefore genetically encoded TetR sponging in our biosensor plasmids by including cassettes containing 0, 10, 20, 40, 80, or 160 TetO binding sites (“sponges”)^12^ (Figure 3D) and performed PEG-osmolality dose-response curves as above (Figure 3G). With DNA-based repressor sponging, P*_opuA_* showed its maximal activation (7.8-fold) at the highest osmolality tested (∼880 mOsm/kg) with 20 TetO sites added to the circuit (P*_opuA_*+20xTetO) (Figure 3G). P_BT0779_ was more sensitive, reaching its maximal activation (16.1-fold) at ∼700 mOsm/kg with 80 TetO binding sites added (P_BT0779_+80xTetO) (Figure 3G). The difference in optimal sponging levels between biosensors likely stemmed from a higher baseline TetR expression by P_BT0779_ that is reflected in its lower baseline levels of GFP observed in dose-response curves (Supplementary Figure 3C). These results suggest that sponging effects are promoter-specific, depending on basal activity and induction dynamics (*V_max_*, *K_m_*, cooperativity) of the native promoter. Importantly, the increased dynamic range with TetR sequestration resulted from higher GFP levels at baseline (∼320 mOsm/kg), while GFP repression at high osmolality remained strong, as predicted (Supplementary Figure 3E,F). Furthermore, sponges did not affect the growth rate or carrying capacity of the P*_opuA_* and P_BT0779_ biosensors compared to wild type cells (Supplementary Figure 3G), suggesting that they impose minimal fitness cost on the cells. Overall, these results demonstrate TetR sequestration as a method to fine-tune biosensor dynamics and establish aTc induction as a rapid 3-day protocol to test repressor sponging without additional cloning, bypassing the ∼3-week process required to construct and test new DNA-based sponge variants.

### Optimized biosensor response is specific, robust to fluctuations in growth rate, reversible, and compatible across Bacteroides

Having developed two osmolality-responsive biosensors that respond to induction with PEG *in vitro,* we next set out to verify the specificity of their response. We therefore grew the P*_opuA_*+20xTetO and P_BT0779_+160xTetO biosensors in gradients of the following ionic and non-ionic osmolytes, and compounds predicted to affect osmotic stress: KCl, NaCl, Na_2_SO_4_, (NH_4_)_2_SO_4_, fructose, lactulose, galactose, sucrose, xylose, sorbitol, glycerol, proline, and betaine. The P*_opuA_*+20xTetO biosensor responded to most osmolytes tested, demonstrating that this biosensor is broadly responsive to increases in environmental osmolality (Figure 3H). In contrast, the P_BT0779_+160xTetO biosensor responded strongly to Na_2_SO_4_ and sucrose, minimally to KCl, NaCl, and (NH_4_)_2_SO_4_, and did not respond to other osmolytes (Figure 3H). This may suggest that the previously uncharacterized BT0779 gene responds specifically to particular causes of osmotic, dehydration, or water structure-related stressors. To ensure that our biosensors do not respond to membrane stress outside of changes to hydration or osmolality, we also tested our P*_opuA_*+10xTetO and P_BT0779_+160xTetO biosensors in increasing concentrations of several membrane-damaging agents (bile salts, Triton-X 100, saponin, and sodium dodecyl sulfate [SDS]), to which we observed no biosensor activation (Supplementary Figure 3H).

Because our inverted biosensor design depends on TetR-mediated repression of GFP, we considered whether changes in growth rate could impact TetR levels and decouple biosensor output from true promoter induction. To explore this, we used our ODE model incorporating TetR sponging to simulate biosensor output across a range of osmolalities and growth rates. Using these simulations, we identified the conditions where the biosensor is predicted to be most sensitive to growth rate effects. This level was around 520 mOsm/kg, where the biosensor should be slightly activated, with repressor levels near the *K_m_* of the repressible promoter. Here, changes in GFP in response to fluctuating TetR concentrations will be the strongest, and lead to increased noise compared to a limited true signal (Supplementary Figure 3I).

To confirm the size of the effect experimentally, we grew the P*_opuA_* and P*_opuA_*+20xTetO biosensors *in vitro* at both baseline and slightly induced conditions where the biosensor is predicted to be most sensitive to growth rate effects (510 mOsm/kg) while modulating growth with graded concentrations of the surfactant SDS which negatively impacts growth rate without inducing an osmotic response (Supplementary Figure 3H). Growth curves were measured in parallel to quantify growth rates at each condition (Supplementary Figure 3J). For each strain and osmolality, signal output was normalized to the baseline condition (base osmolality and 0% SDS) to assess how growth affected the signal. After 6 hours, both biosensors exhibited a consistent ∼2.3-fold ± 0.4 activation at 510 mOsm/kg across normalized growth rates from 0.5–1.0, while showing minimal activation and variation at baseline (0.97-fold ± 0.08) (Supplementary Figure 3J).

Next, to determine whether the strong responses of these biosensors could repeatedly detect and recover from osmotic shifts, a requirement for long-term environmental monitoring, we cycled cultures between baseline and high-osmolality conditions. In both biosensors, activation reversibly increased upon transfer to high osmolality (up to 4-fold for P*_opuA_*+20xTetO and 12-fold for P_BT0779_+160xTetO; Figure 3I), demonstrating that biosensor output accurately tracks environmental changes.

Finally, as other *Bacteroides* spp. can also drive fluorescence from the parent phage promoter^3^, we tested the portability of our biosensor constructs across different hosts. In *B. eggerthii*, *B. finegoldii*, and *B. ovatus*, the P*_opuA_*+20xTetO biosensor showed a graded response to increasing osmolality with a maximum ∼7-, ∼5-, and ∼12-fold activation, respectively (Supplementary Figure 3K). These results suggest that conservation of regulatory elements across this genus may support the function of these fluorescent biosensors in a broader host range.

Altogether, these results show that TetR sequestration improves the dynamic range of the transcriptional reporter circuit and offers a method to fine-tune biosensor activity *in vitro*. The optimized biosensors showed strong, reversible, and osmolality-specific responses, minimal fitness cost, and functional portability across multiple *Bacteroides* species, making them well-suited for further *in vivo* development.

### Repressible promoter variant and GFP degradation tune biosensor dynamics

While repressor sponging offers one strategy to tune biosensor dynamics, another approach involves driving GFP expression from repressible promoters with distinct induction characteristics, thereby modulating the biosensor’s response to repressor levels. To expand the dynamic range and tunability of our biosensors, we constructed variants in which GFP was driven by repressible P_TetR_, P_LacI_, or P_PhlF_ promoters from our libraries, while maintaining repressor expression from the osmolality-sensitive P_BT0779_ promoter. The variants tested exhibited diverse activation profiles across osmolality gradients (Supplementary Figure 4A,B). When tested with and without small molecule inducer, biosensors incorporating P_LacI_ or P_PhlF_ promoters show the highest activation without inducer, suggesting their dynamic range was already optimal without repressor sponging (Supplementary Figure 4A,B). Overall, the behaviour of biosensor variants built with different promoters (including dynamic range, sensitivity, and cooperativity) aligned well with parameters fit in the promoters’ small-molecule induction curves (*V_max_*, *K_m_*) (Figure 1D), confirming that promoter selection alone can be used to systematically tune biosensor performance without circuit redesign.

A limitation of our current circuit design is that GFP turnover depends entirely on cell growth. Without active degradation, TetR-mediated repression reduces GFP only via dilution during cell division, delaying or preventing signal changes when growth is halted. Confirming this, exposing the P*_opuA_*+20xTetO biosensor to high osmolality in minimal media with 0% glucose halted growth and eliminated biosensor activation, while cultures in media with 2% glucose continued to grow and displayed signal turnover (∼2-fold activation; Supplementary Figure 4C). Addressing this, we hypothesized that targeted GFP degradation could enable growth-independent signal change and shorten biosensor response time. We therefore tested whether degradation of GFP could improve biosensor temporal response. Simulations incorporating constitutive GFP degradation showed that while degradation reduced GFP half-life even in non-growing cells (Supplementary Figure 4D,E), it also lowered overall signal and introduced greater sensitivity to growth rate, posing trade-offs for biosensor dynamic range and interpretability (Supplementary Figure 4E-G). Moderate degradation rates offered a balance between signal turnover and baseline expression; we therefore experimentally tagged mGL with the native *Bacteroides* SsrA tag (GETNYALAA)^41,42^. However, this tag caused near-complete signal loss and significantly reduced biosensor activation (∼2-fold vs. ∼7-fold), indicating overly aggressive degradation (Supplementary Figure 4H,I). These results demonstrate that while protein turnover can, in theory, enhance temporal resolution, achieving this requires carefully tuned degradation systems that avoid compromising baseline signal or circuit function. Future work could focus on developing milder, tunable degradation tags to achieve the predicted balance between rapid response and sufficient baseline signal.

### Identification of an alternative integration site enhances P_opuA_ biosensor performance

The broad and consistent *in vitro* response of our P*_opuA_* biosensor to diverse osmolytes (Figure 3H) makes it a promising malabsorption biosensor *in vivo*, where malabsorption can result from a wide range of dietary compounds, pharmaceuticals, or pathologies. To evaluate the performance of our P*_opuA_* biosensors under physiological conditions, we tested their response to increased gut osmolality in a murine model of PEG-induced malabsorption. We first colonized germ-free mice with either the P*_opuA_*+20xTetO biosensor or the NP control and allowed 1-2 weeks for colonization. After equilibration, we treated mice with 0%, 10%, or 15% PEG in drinking water for 6 days. Following treatment, we quantified cecal osmolality as a measure of PEG-induced malabsorption and measured biosensor activation from cecal contents using flow cytometry (Figure 4A; Supplementary Figure 5A,B). At Day 6, mice colonized with either bacterial strain had average cecal osmolalities of ∼380 mOsm/kg without PEG treatment, and significantly increased cecal osmolalities of ∼550 mOsm/kg and ∼650 mOsm/kg in the 10% and 15% PEG treatment groups, respectively (Figure 4B). This result confirmed that PEG administration effectively induces malabsorption and increases gut osmolality. However, despite these increased osmolality levels, both the P*_opuA_*+20xTetO biosensor and the NP control strain showed similar minimal activation levels in all treatment conditions (<1.1-fold activation), when single-cell GFP and RFP levels were measured by flow cytometry (Figure 4C). We reasoned that the lack of P*_opuA_*+20xTetO response *in vivo* could be due to the limited cecal osmolality reached; *in vitro*, the activation of this biosensor is less than half maximal at ∼600 mOsm/kg and continued to increase up to 880 mOsm/kg (Figure 3G). The osmolality reached with 15% PEG treatments *in vivo* may therefore not have been sufficient to trigger a measurable response.

**Figure 4.**
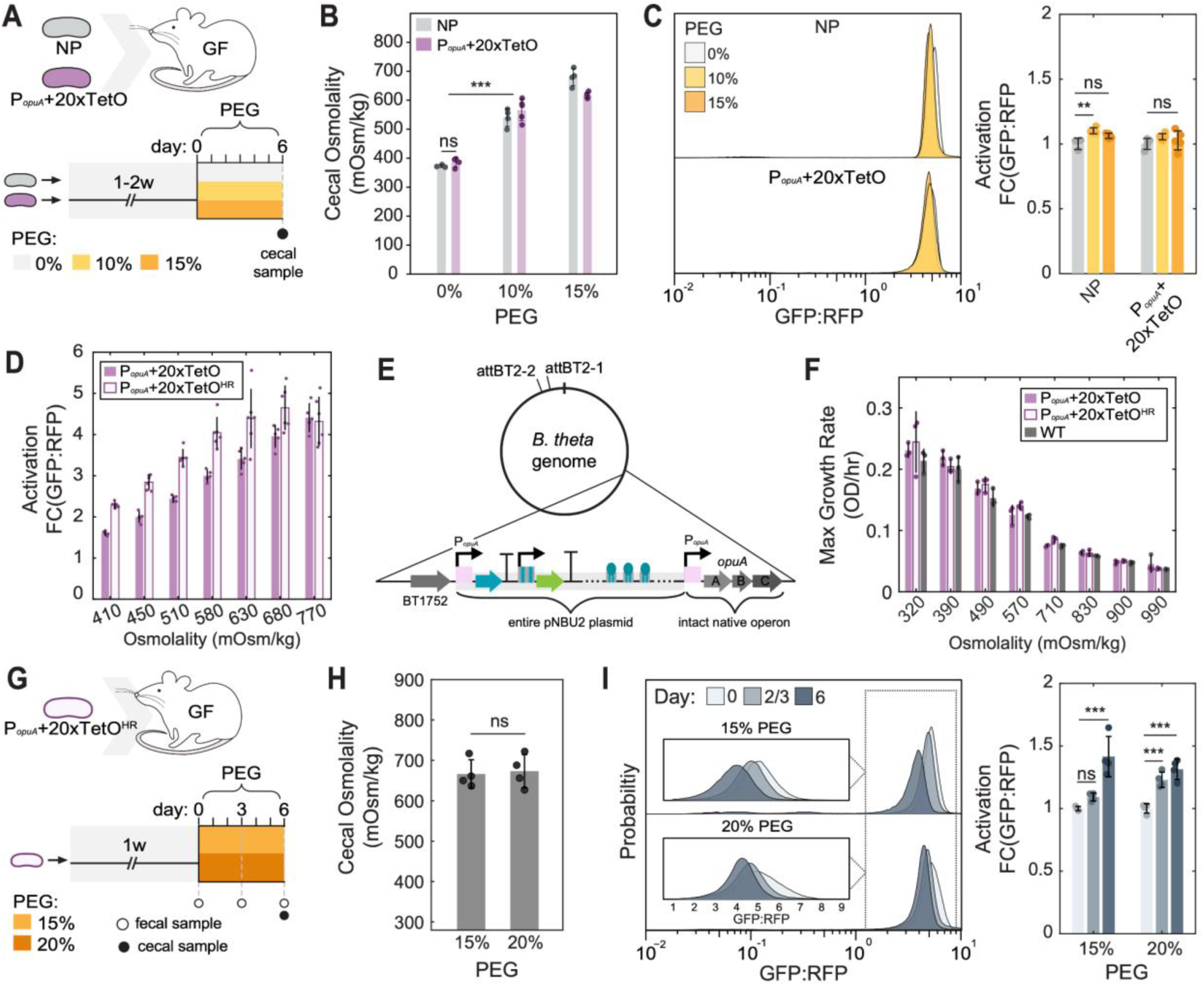
**Integration site of transcriptional reporter circuit enhances *B. theta* osmolality sensing *in vitro* and in a malabsorption mouse model.** A) Experimental outline of PEG-induced malabsorption mouse model. Mice were mono-colonized with either the NP, or P*_opuA_*+20xTetO and treated with 0%, 10%, or 15% PEG in the drinking water for 6 days. B) Cecal osmolality measurements after 6 days of 0%, 10%, or 15% PEG treatment. C) *(left*) Single-cell GFP:RFP ratios measured from cecal contents with flow cytometry for 0%, 10%, and 15% PEG treatments. (*right*) Biosensor activation for each biosensor strain from flow cytometry data. D) Activation in response to increasing osmolality adjusted with PEG, measured in isolated “homologous recombination” (HR) biosensors variants (P*_opuA_*+20xTetO^HR^) and standard attBT2-integrated biosensors (P*_opuA_*+20xTetO). E) Integration of pNBU2 plasmid can occur at the canonical attBT2-1 or attBT2-2 sites or can undergo homologous recombination in the overlapping promoter sequence without disruption of the native gene. F) Growth rates extracted from growth curves of P*_opuA_*+20xTetO and P*_opuA_*+20xTetO^HR^ biosensors, and wild-type *B. theta* in media at increasing osmolality adjusted with PEG indicate that plasmid insertion via homologous recombination does not affect cell fitness. G) Experimental outline of PEG-induced malabsorption mouse model. Mice were mono-colonized with the P*_opuA_*+20xTetO^HR^ biosensor and treated with 15% or 20% PEG in the drinking water for 6 days. H) Cecal osmolality measurements after 6 days of PEG treatment. I) (*left*) Single-cell GFP:RFP ratios measured from fecal contents with flow cytometry for Days 0, 2/3, and 6 of PEG. (*right*) Fold-decrease in the median GFP:RFP ratio for each biosensor strain from flow cytometry data. B,C,H,I: Bar values represent the mean value for *n* = 3 or 4 mice in a treatment group. Error bars represent standard deviation within a group. C,I: Statistics shown are run using the median GFP:RFP values for each mouse in a treatment group. Error bars represent standard deviation within a treatment group. Stats: Pairwise ANOVAs with multiple comparisons (* *padj* < 0.05; ** *padj* < 0.01; *** *padj* < 0.001) D,F: Error bars represent the standard deviation from *n* = 3 biological replicates (unique colonies) of biosensors in each condition.

During previous *in vitro* characterization of the P*_opuA_*+20xTetO biosensor, we noticed a subset of colonies that displayed stronger activation than expected at osmolalities lower than 680 mOsm/kg (∼550–680 mOsm/kg; Figure 4D). Whole-genome sequencing and targeted PCR revealed that the entire biosensor plasmid had integrated directly upstream of the native *opuA* promoter locus, without disrupting the *opuA* operon (Figure 4E; Supplementary Figure 5C). Importantly, integration at this site did not impact strain fitness at any osmolality level in a growth assay, indicating normal *opuA* function in these cells (Figure 4F; Supplementary Figure 5D,E). We termed this variant P*_opuA_*+20xTetO^HR^ as we predicted that integration at this site occurred via homologous recombination (HR), facilitated by ∼500 bp of sequence homology to the *opuA* promoter. Supporting this, we also identified similar variants for the P_BT0779_ biosensor that had integrated at its homologous genomic locus (Supplementary Figure 5C). To determine whether genomic homology increased the likelihood of integration outside the attBT2 sites, we conjugated both the P*_opuA_* biosensor (containing promoter homology) and a P_TetR56_-GFP construct (lacking homology; Figure 1D) and quantified the frequency of insertion sites for 31 colonies of each by PCR. Approximately 19% of P*_opuA_* biosensor colonies lacked attBT2 insertions, compared to only ∼10% of the P_TetR56_-GFP colonies, indicating that homology enhances homologous recombination-based plasmid integration in *B. theta*.

To determine why the HR variant of the P*_opuA_*+20xTetO biosensor shows an enhanced response, we compared its behavior to the original construct integrated at attBT2. Interestingly, the HR variant exhibited consistently lower GFP levels across osmolalities, indicating higher TetR expression, while RFP levels driven by a constitutive promoter remained similar across conditions and strains (Supplementary Figure 5F). These findings imply that promoter activity from P*_opuA_* is influenced by genomic context, potentially due to proximity to native regulatory elements, although further work is needed to confirm this. We conclude that in agreement with studies in *E. coli*^43,44^ and other bacteria^45^, genomic location can impact circuit dynamics in *B. theta*, and that homologous recombination presents a strategy for targeted plasmid integration and potential circuit tuning in *Bacteroides*.

Given that the P*_opuA_*+20xTetO^HR^ biosensor variant showed an increased dynamic range at physiologically osmolalities *in vitro* (<680 mOsm/kg; Figure 4D), we hypothesized that this biosensor could display an improved response in our PEG-malabsorption mouse model. To test this, we colonized germ-free mice with the P*_opuA_*+20xTetO^HR^ strain and treated them with either 15% or 20% PEG in drinking water. During treatment, we collected fecal samples on Days 0, 2 (20% PEG group) or 3 (15% PEG group), and 6 of treatment (Figure 4G), and measured biosensor fluorescence using flow cytometry. On Day 6, mice treated with both 15% and 20% PEG were euthanized and cecal contents were collected for osmolality measurements and flow cytometry analysis. Mice in both treatment groups had an average cecal osmolality of ∼660 mOsm/kg (Figure 4H), comparable to the levels previously obtained with P*_opuA_*+20xTetO (Figure 4B). We observed that the P*_opuA_*+20xTetO^HR^ biosensor showed significant activation by Day 6 of treatment with both 15% and 20% PEG (Figure 4I). Similar final activation levels were reached in the two treatment groups (1.4- and 1.3-fold activation with 15% and 20% PEG treatment, respectively). However, measurements from fecal samples collected at multiple timepoints revealed that the biosensor was activated faster in higher PEG concentrations: in 20% PEG treatment, biosensors reached a significant 1.2-fold activation by Day 2, whereas in 15% PEG treatment, biosensors had not yet responded on Day 3 (Figure 4I). These results confirm that the P*_opuA_*+20xTetO^HR^ effectively detects PEG-induced malabsorption *in vivo* and demonstrate that the genomic integration site plays a crucial role in optimizing biosensor sensitivity and response dynamics. By leveraging targeted integration strategies, this approach enhances the potential for continuous, non-invasive monitoring of the gut environment via fecal sampling.

### An optimized biosensor circuit reports on osmotic perturbations in an *in vivo* mouse model of laxative-induced malabsorption

While our P*_opuA_*+20xTetO^HR^ biosensor effectively responded to the high intestinal osmolalities induced by the consumption of high levels of PEG, a sensor capable of detecting more subtle osmolality shifts would be valuable for diagnosing less severe forms of malabsorption. Given its enhanced sensitivity *in vitro* (Figure 3G), we predicted that our P_BT0779_ biosensor would provide a more sensitive response to PEG-induced malabsorption *in vivo*. Therefore, we tested the response of the biosensor under the high levels of malabsorption tested with the P*_opuA_* biosensors. We colonized germ-free mice with either P_BT0779_+160xTetO or the NP control and allowed 4 weeks for colonization. After equilibration, we treated mice with 0%, 10%, or 15% PEG in drinking water for 6 days to stimulate malabsorption (Figure 5A). Mice treated with 10% and 15% PEG had average cecal osmolalities of ∼550 and 650 mOsm/kg, respectively, with no significant difference in osmolality between mice colonized with NP or P_BT0779_+160xTetO biosensors (Figure 5B). All PEG-treated groups had significantly higher cecal osmolality than untreated mice (∼375 mOsm/kg) (Figure 5B). As expected, the NP control showed no activation after PEG treatment, while P_BT0779_+160xTetO showed a 55.1- and 48.3-fold decrease in GFP:RFP levels in 10% and 15% PEG-treated mice, respectively, compared to untreated mice and the NP control (Figure 5C). The majority of single P_BT0779_+160xTetO biosensor cells in both 10% and 15% PEG-treated mice reached similar GFP:RFP levels to a control strain not producing GFP (“OFF”), suggesting that the biosensor reached maximum induction in these conditions (Figure 5C). As observed *in vitro*, decreasing GFP:RFP levels are the result of decreased GFP levels while RFP levels remained stable (Supplementary Figure 6A). These results indicate that P_BT0779_+160xTetO is highly sensitive to PEG-induced malabsorption in the gut.

**Figure 5.**
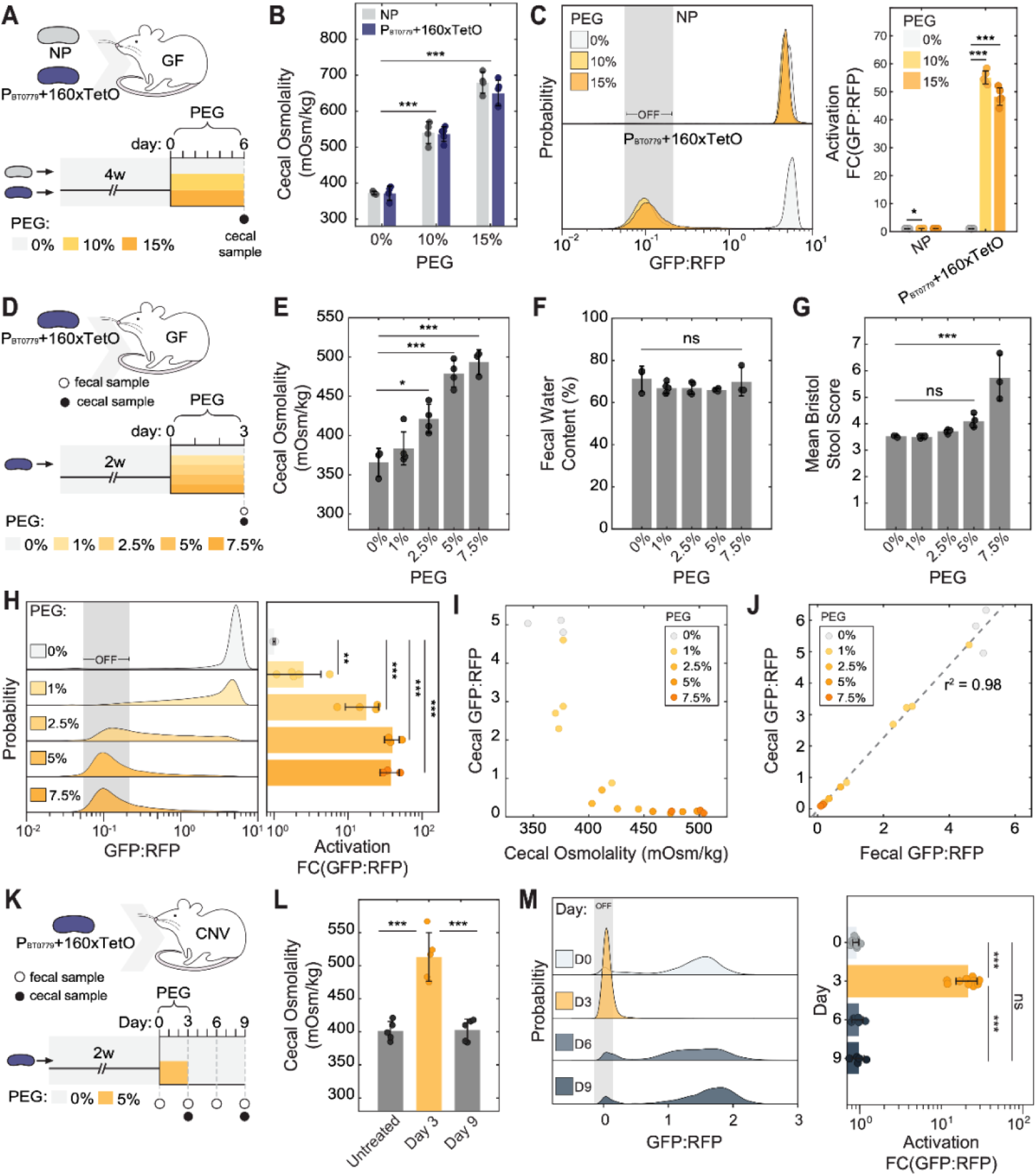
**A *B. theta* biosensor can detect laxative-induced malabsorption at subclinical osmolality levels non-invasively.** A) Outline of initial biosensor test *in vivo*. Mice were mono-colonized with either the NP control or P_BT0779_+160xTetO biosensor, and treated with 0%, 10%, or 15% PEG in the drinking water for 6 days. Cecal contents were collected at D6. B) Cecal osmolality measurements after 6 days of PEG treatment. C) (*left*) Single-cell GFP:RFP ratios measured via flow cytometry from cecal contents of 3-4 mice in each treatment group. Shaded gray area represents background GFP fluorescence levels as measured from a control strain that produces constant RFP but no GFP. (*right*) Fold decrease in the median GFP:RFP ratio for each biosensor strain from flow cytometry data. Dots indicated median GFP:RFP values for each mouse. Statistics are run on median GFP:RFP values. D) Outline of mouse experiment examining the response of P_BT0779_+160xtetO in subclinical osmolality levels. Mice were mono-colonized with P_BT0779_+160xTetO and treated with 0%, 1%, 2.5%, 5%, or 7.5% PEG in the drinking water for 3 days. Cecal and fecal contents were collected after 3 days. E) Cecal osmolality measurements after 3 days of PEG treatment. F) Fecal water content measured^46^ after 3 days of PEG treatment. G) Bristol Stool Scores^47^ from fecal pellets 3 days after PEG treatment. H) (*left*) Single-cell GFP:RFP ratios measured from fecal contents with flow cytometry. (*right*) Fold-decrease in the mean GFP:RFP ratio for each biosensor strain from flow cytometry data. Statistics shown are run on the mean GFP:RFP values for each mouse. I) Median cecal GFP:RFP ratio as a function of cecal osmolality for individual mice in all treatment conditions. J) Median GFP:RFP ratio from fecal and cecal samples. Dotted line shows linear fit, calculated with fit.m function in MATLAB. K) Outline of mouse experiment examining the reversible response of biosensor to osmotic challenge in a complex microbiota context. Conventional mice were colonized with the P_BT0779_+160xTetO, treated with 5% PEG in the drinking water for 3 days, and then allowed to recover for 6 days. Fecal contents were collected prior to PEG treatment (D0), at the end of PEG treatment (D3), and 3- and 6-days post-PEG treatment (D6 and D9, respectively). L) Cecal osmolality measurements for untreated mice, and for mice after 3 days of PEG (D3) and 6 days of recovery (D9). M) (*left*) Single-cell GFP:RFP ratios measured via flow cytometry from fecal contents of 5-10 mice. Shaded gray area represents background GFP fluorescence levels as measured from a control strain that produces constant RFP but no GFP. (*right*) Fold decrease in the median GFP:RFP ratio of biosensor strain from flow cytometry data. Dots indicated median GFP:RFP values for each mouse. Statistics are run on median GFP:RFP values. B, C, E, F, G, H, L, M: Bar values represent the mean value for *n* = 3-10 mice in a treatment group. Error bars represent standard deviation within a group. Stats: Pairwise ANOVAs with multiple comparisons (* *padj* < 0.05; ** *padj* < 0.01; *** *padj* < 0.001)

Next, we investigated whether the P_BT0779_+160xTetO biosensor could detect mild PEG-induced malabsorption by responding to subtle osmolality shifts, even before overt clinical symptoms such as increased fecal water content and decreased stool consistency become apparent. To test this, we colonized germ-free mice with P_BT0779_+160xTetO and treated them with either 0%, 1%, 2.5%, 5%, or 7.5% PEG for 3 days, after which we collected cecal and fecal samples (Figure 5D). Mean cecal osmolalities reached 366, 384, 422, 479, and 493 mOsm/kg for the 0%, 1%, 2.5%, 5%, or 7.5% treatment groups, respectively (Figure 5E). While cecal osmolality increased gradually with PEG treatment, standard non-invasive measures of malabsorption, such as fecal water content^46^ and feces consistency as measured by Bristol Stool Scoring^47^, did not differ between treated and untreated mice, except for 7.5% PEG (Figure 5F,G). The level of malabsorption present would therefore not be detectable by standard clinical methods. The P_BT0779_+160xTetO biosensor showed strong activation in response to increasing osmolality, measured from both fecal (Figure 5H) and cecal samples (Figure 5I,J). Similar to previous experiments, changes in GFP:RFP ratio are the result of decreased GFP levels while RFP levels remained stable (Supplementary Figure 6B). Biosensors from fecal samples were most activated in mice treated with 5-7.5% PEG, showing an average ∼40-fold activation compared to untreated mice (Figure 5H). In individual mice, P_BT0779_+160xTetO GFP:RFP ratios from cecal samples correlate well with cecal osmolality (Figure 5I), showing this sensor’s ability to discern even small differences in osmolality. Notably, GFP:RFP measurements from fecal and cecal samples were strongly linearly correlated (r^2^ = 0.98; Figure 5J), demonstrating that biosensor response can be reliably measured non-invasively from stool samples.

To verify the suitability of our biosensor for long-term monitoring in the context of a complex gut microbiota, we administered a single dose of the P_BT0779_+160xTetO biosensor to conventional mice and tracked its behaviour over time. First, to assess the stability of our biosensor in a conventional microbiota, we tracked biosensor colonization and colony fluorescence over four weeks. Colonization stabilized at ∼10^5^-10^6^ CFU/mL feces within 7 days (Supplementary Figure 6C), consistent with prior studies of engineered *B. theta* in complex communities^4,10^. Fluorescent signals remained stable over time, with >90% of colonies on average retaining both GFP and RFP by 31 days of colonization, suggesting that *in vivo* fluorophore expression does not strongly impact fitness and that loss of fluorescence is not under positive selection (Supplementary Figure 6D). Further supporting this, the ratio of GFP^+^ to RFP^+^ colonies in individual mice shifted dynamically over time, with some timepoints showing a predominance of GFP^+^ colonies and others of RFP^+^ colonies (Supplementary Figure 6E). This variability suggests that reporter mutations did not become fixed in the population, supporting the long-term stability of reporter expression in a complex gut environment.

Next, to verify that our biosensor signal is transient and recoverable, we measured biosensor response before, during, and after treatment with 5% PEG using flow cytometry (Figure 5K; Supplementary Figure 6F-H). By Day 3 of PEG treatment, cecal osmolality had increased from ∼400 to ∼500 mOsm/kg (Figure 5L), and the biosensor showed ∼21-fold activation (Figure 5M). Importantly, both cecal osmolality and biosensor signal returned to baseline levels within three days post-treatment and remained stable for at least three more days (Figure 5L,M). As expected, observed signal changes were due to changing GFP levels, while RFP levels remained stable (Supplementary Figure 6F,H). In both this experiment and previous mono-colonization experiments, baseline GFP levels in gated RFP^+^ cells remained stable across all periods of experiments without PEG treatment (Supplementary Figure 6I), demonstrating its long-term stability in the gut. Together, these results demonstrate that the P_BT0779_+160xTetO biosensor functions robustly and reversibly across diverse community contexts, reliably detecting osmolality shifts *in vivo* from cecal and fecal samples. Importantly, the biosensor is sensitive enough to differentiate malabsorption severity prior to overt clinical symptoms, supporting its utility for precise, non-invasive, non-invasive, long-term gut environmental monitoring.

Taken together, our results provide novel synthetic biology tools for *B. theta* engineering, identify osmolality-responsive promoters, and validate biosensors capable of detecting and reporting malabsorption of relevant pharmaceutical and dietary compounds *in vitro* and *in vivo*. These findings demonstrate the potential of engineered *B. theta* for monitoring of gut physiological states and support the potential impact of microbiota-based diagnostics for gut health.

## Discussion

In this work, we developed and validated two *B. theta* biosensors that robustly detect environmental osmolality shifts both *in vitro* and in laxative-induced malabsorption models. These biosensors enable sensitive, continuous, and non-invasive monitoring via fecal sampling, demonstrating their potential for real-time gut diagnostics. Their development was made possible by the expansion of the *B. theta* synthetic biology toolkit, to which we contributed a library of novel small-molecule-inducible promoters, a DNA-based repressor sponging system adapted from *E. coli*, and a modular fluorescence-based transcriptional reporting circuit. Additionally, we characterized a novel synthetic plasmid insertion mode in *B. theta,* providing insights into how genomic integration site can influence biosensor performance. Beyond the applications presented here, our fluorescence-based *B. theta* genetic tools expand the available parts for building complex synthetic circuits in this commensal bacterium. These advancements not only expand our ability to study *B. theta* biology, but also strengthens its potential as a chassis for microbiota-based diagnostics.

For effective long-term, continuous monitoring applications, microbial-based biosensors must achieve stable and abundant gut colonization in addition to detecting and responding to environmental changes in the gut. Historically, *E. coli* has been the preferred chassis for gut bacterial biosensors due to its expansive genetic toolkit and ease of engineering^48,49^. For example, biosensors engineered in probiotic *E. coli* Nissle 1917 and murine commensal *E. coli* NGF-1 strains have been used to detect and report on inflammation^50–53^ and colorectal tumors^54,55^ in the GI tract. Despite these developments, even probiotic *E. coli* strains can only colonize transiently and sparsely following disruption of the native microbiota with antibiotics or intestinal lavage^56,57^. This limits long-term applications for these biosensors, and risks reporting from a perturbed gut environment. In contrast, *B. theta* is a prevalent and abundant bacterium in the human microbiota, with ∼46% prevalence in human microbiota^58^ and composing ∼6% of the bacterial microbiota when present^59^. Engineered *B. theta* strains can stably colonize an existing conventional microbiota in mice and humans^4,10^, circumventing the need to perturb the native microbiota. Members of the *Bacteroidaceae* family also naturally colonize across gut regions^60^ and in diverse niches of the colon (e.g., the lumen and outer mucus layer^61^, and the intestinal crypts^5,62^, an important feature for detecting divergent responses from spatially distinct populations. Finally, engineering *B. theta* to degrade the exogenous metabolite porphyrin was recently shown to allow tuneable levels of colonization in the gut^4,10^, which could give the user control over the survival and abundance of engineered strains. Together, its stable colonization, broad niche occupancy, and potential for tunable population control within the gut environment make *B. theta* a compelling chassis for gut biosensors and engineered genetic circuits.

The ability of the biosensors developed here to produce fluorescence outputs expands the possibilities for multiplexing reporters. Unlike previously used luminescence outputs, the distinct excitation and emission profiles of fluorescent proteins allows reporter signal detection from multiple fluorophores. Combined with the orthogonality of the three repressible promoter libraries presented here, utilizing fluorescence output could therefore enable the construction of multiplexed reporters to process multiple inputs simultaneously. For example, combining the P_BT0779_+160xTetO and P*_opuA_*+20xTetO^HR^ biosensors presented here could combine the sensitivity and robustness of each circuit to form a system capable of detecting and differentiating causes and severity of malabsorption. Combining malabsorption detection with other triggers could also be useful to differentiate gut pathologies with overlapping symptoms based on the gut environment. For example, biosensors that detect both malabsorption and inflammation-related perturbations (e.g., increased levels of reactive oxygen species^63^ or nitrate^9^) could aid in differentiating gut pathologies such as celiac disease and IBD that present both of these symptoms, from other dietary intolerances that cause malabsorption alone (e.g., lactose, sorbitol, or fructose intolerances)^15^. Rapid assembly and characterization of such multiplexed biosensors is facilitated by our modular biosensor backbone and parts vectors. Furthermore, our transcriptomics analysis of *B. theta* under different environmental conditions already suggests candidate promoters that could be integrated to make biosensors for a range of environmental triggers (Figure 2C). On a large scale, this system could also be used to assemble a comprehensive reporter library for *B. theta*, which could help characterize this bacterium’s response and adaptation to diverse conditions in the gut and identify promoters that respond to clinically relevant environmental cues.

Single-cell transcriptional analyses have become a powerful tool for understanding gut biogeography, with advances in droplet-based^64,65^, combinatorial barcoding^66–68^, fluorescence *in situ* hybridization (FISH)^11^, and transcriptional recording^69,70^ methods enabling high-throughput, multiplexed screening that broadly characterizes transcriptional responses to gut environment changes. However, these approaches are experimentally demanding, require extensive sample processing, and/or rely on deep sequencing, limiting their application in rapid biosensing for gut monitoring. Additionally, their untargeted nature can bias detection toward highly expressed genes, potentially overlooking small but environmentally relevant transcriptional changes. In contrast, synthetic biosensing circuits use specific promoters to selectively detect gene activity changes, allowing for targeted measurement of specific responses—even for lowly expressed genes overlooked by untargeted approaches. Furthermore, fluorescence reporters facilitate rapid, single-cell analysis via flow cytometry with minimal sample preparation. For example, measurement of signal from our *B. theta* biosensors took ∼3 hours for up to 24 samples, from fecal collection to completion of flow cytometry measurements. Processing time was often limited by the maturation time of the constitutive RFP in our circuit (∼2 hours), rather than experimental procedures. In situations where rapid processing is imperative and sensitive reporting of a limited number of transcripts is sufficient (e.g., detection of limited factors for diagnostics), our synthetic biosensor approach allows near real-time signal detection and non-invasive sampling, therefore offering a streamlined option for transcriptional analysis.

The transcriptional reporter system and resulting biosensors presented here can help us interrogate important questions about the gut microbiota and its biogeography in health and disease. The malabsorption detectors characterized in this system could be used to characterize osmotic changes to the gut environment in clinically relevant models affecting malabsorption, intestinal permeability, or hydration, or such as pathogen invasion (e.g., *Trichinella* spiralis^71^ or enteropathogenic *E. coli* infection^72^), malabsorptive conditions (e.g., Celiac disease^73^, dietary perturbations such as high-fat diet^74^, or inflammatory pathologies (e.g., dextran sodium sulfate^75^ or immunodeficiency^76^ colitis models). Moving towards diagnostics in the human microbiome, gnotobiotic mouse models or gut-on-chip platforms could facilitate the translation of our biosensors to a diagnostic setting without direct administration of these engineered microbes. Along with the novel synthetic biology tools characterized in this work, these tools support the use of *B. theta* in gut environmental monitoring and point to the utility of commensal gut bacteria as diagnostic and therapeutic tools for gut health.

## Limitations

While our biosensors perform robustly in diverse gut environments, they are subject to several biological and technical limitations. Fluorescent protein expression in *Bacteroides* enables single-cell resolution via FACS and endpoint microscopy^3,4,77^; however, as fluorophore maturation requires oxygen, use for live imaging within the anaerobic gut is restricted for these sensors. In such cases, luminescence reporters would be preferable and have been successfully applied for live gut imaging of other model bacteria^78,79^.

Another limitation of the biosensors presented here lies in the use of an inverted reporter logic, where biosensor activation depends on repression of GFP and its dilution upon bacterial growth. Under no-growth conditions, output therefore remains unchanged even if the stimulus is present (Supplementary Figure 4C). Furthermore, variations in growth rate can lead to TetR accumulation or altered GFP dilution, affecting signal output independently of promoter activity. Results from control experiments altering growth at a given induction level (Supplementary Figure 3I,J) suggest that growth-related effects were not the main driver of biosensor signal change and lead to a <1.4-fold variation in activation. Nonetheless, this effect introduces a non-linear relationship between stimulus strength and signal amplitude that may complicate interpretation, especially in the gut, where bacterial growth dynamics remain poorly characterized.

Promoter selection also posed challenges. Several candidates did not elicit a measurable response (Figure 3B), likely due to weak activity or improper regulation: given the under-characterization of *B. theta* promoter architecture, key regulatory motifs may have been inadvertently omitted during biosensor construction. Refining transcriptomic analysis to prioritize absolute expression levels over fold change could aid in selecting stronger promoters. Additional circuit improvements—such as introducing GFP degradation tags or using different repressible promoters (Supplementary Figure 4)—could further enhance dynamic range and sensitivity.

Finally, biosensor performance *in vivo* can be affected by evolutionary and colonization dynamics. Mutations in the reporter circuit that become fixed may generate false positives or negatives. Moreover, some microbiotas may resist stable *B. theta* engraftment^4^. Engineering approaches that promote *B. theta* colonization^4,10^ could help mitigate these limitations.

## Conclusion

By expanding the synthetic biology toolkit for *B. theta* and demonstrating its application as a biosensor for gut osmolality, this work advances the potential of sensitive microbiota-based diagnostics for real-time, non-invasive gut health monitoring. Beyond malabsorption, the tools and approaches developed here provide a foundation for engineering biosensors to detect the activation of *B. theta* promoters as well as a wide range of gut conditions, including inflammation, pH shifts, and metabolite fluctuations. These innovations establish a foundation for using engineered *Bacteroides* in non-invasive diagnostics and microbiome-based therapeutics.

## Resource availability

### Lead contact

Requests for further information and resources should be directed to and will be fulfilled by the lead contact, Carolina Tropini

### Materials availability

All plasmids containing characterized promoters, TetR sponge cassettes, biosensor entry vectors, and assembled biosensor circuits generated in this study (Supplementary Table 6) have been made available on Addgene.

## Methods

### Animal Husbandry

Animal experiments at The University of British Columbia were performed in accordance with The University of British Columbia Animal Care Committee. Gnotobiotic mice were generated from germ-free (GF) Swiss Webster colonies. Gnotobiotic and conventional Swiss Webster mice were maintained in gnotobiotic isolators or Ehret cages, respectively. For experiments, mice were transferred and housed in ISOcage P-Bioexclusion System (Tecniplast) under constant high positive pressure and fed an autoclaved standard diet (Purina Lab Diet 5K67) for the duration of the experiments. Mice were sacrificed at indicated time points using carbon dioxide asphyxiation, followed by secondary euthanasia of cervical dislocation.

### Bacterial strains and culture conditions

All *E. coli* strains used in this study were grown aerobically at 37 °C in LB Miller broth (Fisher BP1426500) or LB Miller plates with 1.5% agar (Fisher BP9744500). Where appropriate, antibiotics were added at the following concentrations: ampicillin/carbenicillin at 100 μg/mL (Fisher BP1760-25/Fisher BP26485); kanamycin at 50 μg/mL (Sigma Aldrich K1377); chloramphenicol at 25 μg/mL (Acros Organics AC227920250). Wild type *B. theta* VPI-5482 (ATCC 29148; GenBank AE015928.1) was used as a base strain for all genetic engineering in *B. theta*. Other *Bacteroides* engineered include *B. caccae* (ATCC 43185); *B. cellulosilyticus* (DSMZ 14838); *B. distasonis* (ATCC 8503); *B. eggerthii* (ATCC 27754); *B. finegoldii* (CL09T03C10 BEI HM-727); *B. fragilis* (NCTC 9343); *B. ovatus* (ATCC 8483); *B. salyersiae* (CL02T12C01 BEI HM-725); *B. uniformis* (ATCC 8492); and *B. xylanoisolvens* (DSMZ 18836). All *Bacteroides* strains were grown in either Brain-Heart Infusion broth (BD B11059) supplemented with 0.5 μg/mL vitamin K1 (Alfa Aesar AAL1057506) and 5 μg/mL hemin chloride (MilliporeSigma 37415GM) (BHIS) or in Tryptone-Yeast extract-Glucose media (10 g/L Tryptone [Gibco 211705]; 5 g/L Yeast Extract [Gibco 212750]; 2 g/L glucose [Fisher BP350500]; 1 M combined K_2_HPO_4_ and HK_2_O_4_P, pH 7.2 [Fisher BP362-1 and Fisher BP363-1 combined to pH 7.2], 0.8% CaCl_2_ [Fisher BP510-500]; 0.4 mg/mL FeSO_4_ [Alfa Aesar AAA1517836]; 0.02 g/L MgSO_4_ [Fisher BP213-1]; 0.4 g/L NaHCO_3_ [Fisher BP328-500]; 0.08 g/L NaCl [Fisher S271]) supplemented with vitamin K1 (0.5 μg/mL), hemin chloride (5 μg/mL) and cysteine hydrochloride (0.5 g/L; BP376-100 Fisher) (TYG). All *Bacteroides* strains were streaked out in BHIS plates with 1.5% agar. Where appropriate, antibiotics were added: gentamicin (Gen) at 200 μg/mL (Sigma Aldrich G3632-5G); erythromycin (Erm) at 25 μg/mL (Alfa Aesar, AAJ6227909). *B. theta* strains were grown in an anaerobic chamber (Mandel) under the following atmospheric conditions: 5% CO_2_ and 2.4-5% H_2_ in N_2_ (Gas Mix: Linde BioBlend NI CD5H11U−T). All plates and media used to grow *Bacteroides* were pre-reduced under anaerobic conditions for at least 12 h before use. All bacterial cell cultures were stored in 25% glycerol (Fisher G331) at −80 °C for long-term storage.

### Genetic parts, plasmid, and strain construction

All genetic parts, primers, and plasmids used in the construction of strains are listed in Supplementary Table 7. All assembled plasmids were sequence verified by Oxford Nanopore sequencing (Plasmidsaurus) and have been deposited on Addgene under the accession numbers specific in Supplementary Table 6.

#### Competent cells and transformations

All cloning and plasmid construction was completed using either *E. coli* S17-1 λpir (ATCC-BAA-2428) or commercially available competent *E. coli* OneShot^TM^ PIR1 cells (Invitrogen C101010). *E. coli* S17-1 λpir were made electrocompetent or chemically competent using the TSS method^80^.

### TSS chemically competent cells

An overnight culture of *E. coli* S17-1 λpir was sub-cultured at a 1:100 ratio and grown ∼2-3 h until an OD_600_ of 0.4-0.5. Cells were then harvested by centrifugation at 3000 x *g* for 10 min in a 4 °C rotor. Supernatant was removed and cells were resuspended in 10% of the original culture volume of TSS buffer (10% w/v PEG 3350 [Sigma 202444]; 0.03 M MgCl_2_ [Fisher BP214]; 5% v/v DMSO [Invitrogen D12345] in LB-Miller broth). Cells were dispensed into 100 μL aliquots and stored at −80 °C until use.

For transformation, DNA was added to 100 μL thawed cells and the mixture was incubated on ice for 30 min. Cells were placed at 42 °C for 30 s and placed back on ice for 2 min. 900 μL SOC recovery media (MP Biomedicals MP113031012) was added, and cells were incubated at 37 °C while shaking at 225 RPM for 1hr. 100-300 μL of transformed cells were then plated on LB-agar plates with appropriate antibiotic selection and incubated at 37 °C overnight. For DNA assemblies, 5 μL of assembly reaction was used for transformation. For all other transformations, 1 μL purified plasmid was added.

### Electrocompetent cells

Overnight cultures of *E. coli* S17-1 λpir were diluted at 1:100 into Erlenmeyer flasks containing fresh LB broth and incubated for ∼3 h until they reached OD_600_ of 0.6-1.0. The cells were incubated on ice for 30 min. Cells were harvested by centrifugation for 15 min at 2300 x *g* in a 4 °C rotor, supernatant was carefully removed, and cells were resuspended gently in ice-cold sterile water. This process was repeated a second time. Cells were centrifuged, supernatant was removed, and cells were resuspended in ice-cold 10% glycerol solution. Cells were transferred into cold centrifuge tubes and incubated on ice for 30 min. Cells were centrifuged again, supernatant was removed, and cells were resuspended in ice-cold 10% glycerol solution. Cells were flash-frozen in liquid N_2_ in 100 µL aliquots and stored at −80 °C.

For transformation, up to 2.5 μL of purified plasmid or 1 μL of Golden Gate assembly reaction was added to 25 µL of thawed cells and the mixture was incubated on ice for 10 min. Cells and DNA were transferred to a cold 2 mm gap electroporation cuvette (BioRad 1652082) and electroporated using a BioRad Gene Pulser Xcell electroporation system at the following settings: Exponential decay pulse, 25 μF, 200 Ω, 2500 V. SOC recovery media was added immediately after electroporation to a volume of 1 mL, and transformed cells were incubated at 37 °C for 1 hr, shaking at 225 RPM. 100-300 μL of transformation was plated onto selective LB-agar plates.

### Construction genetic parts, plasmids, and strains

#### Repressible P_BFP1E6_ promoter libraries

Promoter libraries containing TetR, LacI, or PhlF binding sites were ordered as pools of single-stranded oligonucleotides (IDT oPools). All unique promoter sequences were flanked by universal sequences containing primer binding sites for amplification, BsaI recognition sites, and overhang sequences for assembly into the reporter vector. Promoter parts were prepared for Golden Gate assembly by second strand synthesis: briefly, single-stranded oligonucleotide libraries were resuspended in nuclease-free water (Fisher BP2484100) and annealed with a universal reverse primer in 1X Phusion HF PCR buffer (NEB B0518). Annealed oligonucleotides were then mixed with Phusion® High-Fidelity DNA Polymerase (NEB M0530) and dNTPs (NEB N0447) and extended at 72 °C. Synthesized double-stranded DNA was purified in nuclease-free water using a QIAquick PCR purification kit (Qiagen 28106) following manufacturer protocols.

Existing repressible promoters P1T_DP_ and P_LacO23_ were flanked by BsaI sites and overhang sequences for assembly into the reporter vector, were synthesized using assembly PCR (aPCR). Briefly, two long (∼80-100 bp) oligonucleotides with a 58 °C Tm overlap were designed containing the RBS sequence flanked by BsaI recognition sites at each end. Two short outer primers (∼58 °C Tm) were designed against the 5’ and 3’ end of the longer oligos. Inner and outer primers were mixed at a 1:10 molar ratio and subjected to rounds of amplification following the manufacturer-recommended PCR protocol for Phusion® High-Fidelity DNA Polymerase. PCR products were purified in nuclease-free water using a QIAquick PCR Purification Kit.

#### RBS parts

The strong *B. theta* RBS1P-LP sequence^3^ was constructed as a Golden Gate assembly part using assembly PCR, as above.

#### Repressor parts vectors

TetR and LacI repressor coding sequences without BsaI recognition sites were ordered as synthetic gBlocks (IDT). A Golden Gate-compatible PhlF gene was obtained (pR-PhlF^81^; Addgene 49367). Golden Gate parts vectors containing these repressors under *B. theta* promoters were assembled using HiFi DNA assembly using NEBuilder® HiFi DNA Assembly Master Mix (NEB E2621S). Briefly, a ColE1 plasmid backbone^82^, *B. theta* promoter (PBfP2E5) and RBS (RBS1P-LP)^3^, and repressor genes with a terminator were amplified from existing plasmids using Phusion DNA polymerase and assembled using HiFi DNA assembly. BsaI recognition sites were inserted upstream and downstream of the *B. theta* promoter and terminator, respectively, such that the whole expression cassette can be cut out and assembled into a new plasmid as part of a Golden Gate Assembly. After cloning, parts vectors were purified by miniprep (Qiagen 27106) and verified by Oxford Nanopore sequencing (Plasmidsaurus).

#### Golden Gate-compatible pNBU2 entry vector for repressible promoter screening

Repressor expression cassettes, promoters, and RBS parts described above were assembled into the pNBU2 backbone, which was transferred from *E. coli* to *B. theta* during conjugation and subsequently integrated into the *B. theta* genome in single copy at one of two attBT2 sites. To build a Golden Gate entry vector for osmolality promoters, an existing pNBU2 plasmid (pWW3536^3^ [Genbank KY776518]) containing an *sfgfp* gene was adapted for our purposes. In this plasmid, the constitutive promoter driving *sfgfp* was replaced with an mCherry dropout cassette flanked by BsaI recognition sites. The *mCherry* gene and its terminator were PCR amplified from the pWW3515 plasmid^3^ (Genbank KY776515), with the forward primer containing an overhang coding a strong *E. coli* promoter (J23100) and RBS^82^. Overhangs also contained BsaI recognition sites. The pNBU2 backbone containing the *sfgfp* gene and its terminator was amplified from the pWW3536 plasmid^3^, with primers containing overhangs that overlap the amplified mCherry cassette. PCR products were purified and assembled using HiFi DNA assembly to build an mCherry dropout entry vector, into which parts are assembled into pNBU2 as follows: repressor cassette:BsaI-flanked promoter entry site:RB:GFP:pNBU2 backbone (pEM124; Figure 1B), and colonies containing assembled and background plasmids can be easily distinguished.

#### Promoter library construction in E. coli

Repressors, cognate promoters, and RBS1P-LP were inserted into the pEM124 pNBU2-GFP backbone via BsaI Golden Gate assembly. Assemblies were transformed into *E. coli* OneShot^TM^ PIR1 cells (Invitrogen C101010). ∼4000 colonies were scraped, resuspended into 10 mL selective LB broth, and stored as glycerol stocks in 200 μL aliquots. To transfer the plasmid library to *E. coli* S17-1 λpir cells for conjugations, one 200 μL aliquot was thawed on ice, diluted 1:100 into selective LB broth, grown for 4 h at 37 °C at 225 RPM, and miniprepped. Purified plasmids were transformed into electrocompetent S17-1 𝜆pir cells and plated on selective LB-agar plates. ∼4000 colonies were scraped into 5 mL of selective LB broth and stored at −80 °C in 200 μL aliquots (pEM252) to be used for conjugation.

#### Construction of Golden Gate-compatible biosensor entry vectors

The inverted transcriptional circuit was assembled into the pNBU2 backbone using HiFi assembly as follows: P_PBFP1E6_:RBS:mScarlet-I3:BsaI-flanked entry site:TetR:P_TetR56_:RBS:mGL:pNBU2 backbone (pEM526; Figure 3A). pDx_mScarlet-I3 was a gift from Dorus Gadella (Addgene plasmid 189757). The mGL gene was ordered as a gBlock (Twist Biosciences) and PCR amplified prior to HiFi assembly. Assemblies were transformed into TSS competent *E. coli* S17-1 λpir cells.

#### Construction of biosensor circuits

TetR sponging cassettes were adapted for use with Golden Gate assembly into our circuit. Briefly, plasmids containing 10, 20, 40, 80, or 160 TetO binding sites (Addgene 160825-160829) were digested with XbaI (NEB R0145) and XhoI (NEB R0146) according to manufacturer-recommended protocols. The p15A ori backbone of these plasmids was PCR amplified with primers overlapping sponge cassettes and containing BsaI sites matching the pEM526 entry vector at the upstream end and the native promoter at the downstream end. Sponge cassettes were assembled into these modified backbones with HiFi assembly to make Golden Gate-ready TetR sponging cassettes (EM413, EM414, EM415, EM351, EM352) with overhangs matching the pEM526 entry vector (5’-CCGA-3’) at the 5’ end and a native promoter part (5’-ATCA-3’) at the 3’ end. Native promoters and RBSs were amplified from the *B. theta* genome (∼500 bp upstream of the CDS), with primers containing BsaI sites and overhangs matching either the pEM526 entry vector (5’-CCGA-3’) or sponge cassettes (5’-TGAT-3’) at the 5’ end and TetR (5’-ACAT-3’) at the 3’ end. Promoter and TetR sponge parts were assembled into pEM526 in 10 μL Golden Gate reactions containing:

0.33 μL BsaI HFv2 (NEB R3733G)

0.33 μL T4 DNA Ligase (NEB M0202M)

1 μL 10x T4 DNA Ligase Buffer (NEB B0202S)

25 fmol Entry Vector Plasmid

50 fmol each additional part

To volume with Nuclease Free water (Millipore Sigma 7732-18-5)

Reactions were cycled as follows:

37 °C for 10 min

25x:

37 °C for 1.5 min

16 °C for 3 min

37 °C for 5 min

80 °C for 10 min

12 °C hold

All assemblies were transformed into TSS competent *E. coli* S17-1 λpir cells. Final biosensor plasmids were conjugated into *B. theta* as below.

#### Alternative repressible promoters

Biosensor constructs containing promoters other than P_TetR56_ (Supplementary Figure 4A,B) were assembled in 4-piece HiFi assemblies. Repressible promoters containing overhangs for HiFi assembly, were ordered as gBlocks from Twist Bioscience and amplified with Phusion PCR. The P_BT0779_ biosensor backbone was amplified in two pieces with Phusion PCR, and repressor parts were amplified with overhangs for HiFi assembly from the corresponding Golden Gate parts vectors. Primers for HiFi pieces are listed in Supplementary Table 7.

#### SsrA tagged GFP constructs

A version of the mGL biosensor entry vector was assembled in a 2-piece HiFi assembly. The existing entry vector was linearized via PCR amplification. Overlapping oligonucleotides containing a short linker (GS: 5’-GGATCT-3’), the *Bacteroides* SsrA tag sequence (GETNYALAA: 5’-GGTGAAACCAATTATGCTCTTGCCGCC-3’), a stop codon, and homology to the linearized backbone were amplified with assembly PCR as above and prepared for HiFi Assembly by PCR purification. The tagged entry vector was used to test degradation levels under constitutive GFP expression (NP::SsrA). To build the P*_opuA_*+20xTetO::SsrA construct, the P*_opuA_* promoter and 20-sponge cassette were inserted into the SsrA-tagged entry vector as described above for standard biosensor assemble. Primers for HiFi pieces are listed in Supplementary Table 7.

#### Bacteroides conjugation and integration

Wild type *Bacteroides* strains were grown overnight in BHIS broth in anaerobic conditions, then sub-cultured in 5 mL fresh BHIS at 1:50 dilution. *E. coli* S17-1 λpir containing assembled pNBU2 plasmids were grown overnight in LB-Amp, then sub-cultured in 5 mL fresh LB-Amp. Subcultures for both strains were grown ∼4-5 h, centrifuged at 3000 x *g* for 10 min and supernatant was removed. *E. coli* and *Bacteroides* pellets were serially resuspended in 1 mL of BHIS. 100 μL of resuspension was spot-plated onto BHI+10% Horse Blood (Dalynn Biologicals HH30-500) agar plates and incubated at 37 °C aerobically for 14-16 h, followed by anaerobic incubation at 37 °C for another 14-16 h. Resulting growth from conjugation mixture was spread onto selective BHIS agar plates with gentamicin and erythromycin and grown anaerobically for two nights. Single colonies were re-streaked onto BHIS plates with gentamicin and erythromycin and grown anaerobically for two nights. For *B. theta*, the genomic insertion site of the plasmid was verified in single colonies by colony PCR (*PCR verification of integration site*) before long-term storage. For other *Bacteroides*, integration site was not verified. Single colonies were picked into liquid BHIS + Erm + Gent and grown overnight. Liquid cultures were mixed at 1:1 ratio with 50% glycerol for long-term storage.

In the case of promoter library conjugations, all single colonies from the first selective BHIS plates (4 plates with ∼1000 colonies/plate) were scraped into 10 mL liquid BHIS-Erm-Gen and incubated for 4 h to select against any remaining donor *E. coli*. This liquid culture was mixed at 1:1 ratio with 50% glycerol. To determine the number of colony-forming units in the glycerol stocks, an aliquot of this mixture was diluted at 1:100, 1:1000, and 1:10000 ratios in BHIS, 100 μL of each dilution was spread onto BHIS-Erm plates and grown overnight, and colonies were counted. The remaining glycerol-culture solution was stored at − 80 °C as 100 μL aliquots.

#### PCR verification of integration site

Briefly, a set of primers was designed flanking the attBT2-1 and attBT2-2 insertion sites, with one primer in the pNBU2 backbone such that two bands of specific sizes are produced according to insertion site (Table 1; Supplementary Figure 5C). Colony PCRs were performed with DreamTaq PCR Master Mix (ThermoFisher K1081) according to manufacturer recommended protocols with a total of 1 µM of primers. In the case that biosensor colonies resulting from conjugations produced WT amplicons at both attBT2 sites, integration at the native promoter site was verified using two sets of primers targeting the 5’ and 3’ ends of the plasmid and the flanking genomic regions (Table 2; Supplementary Figure 5C).

**Table 1.**
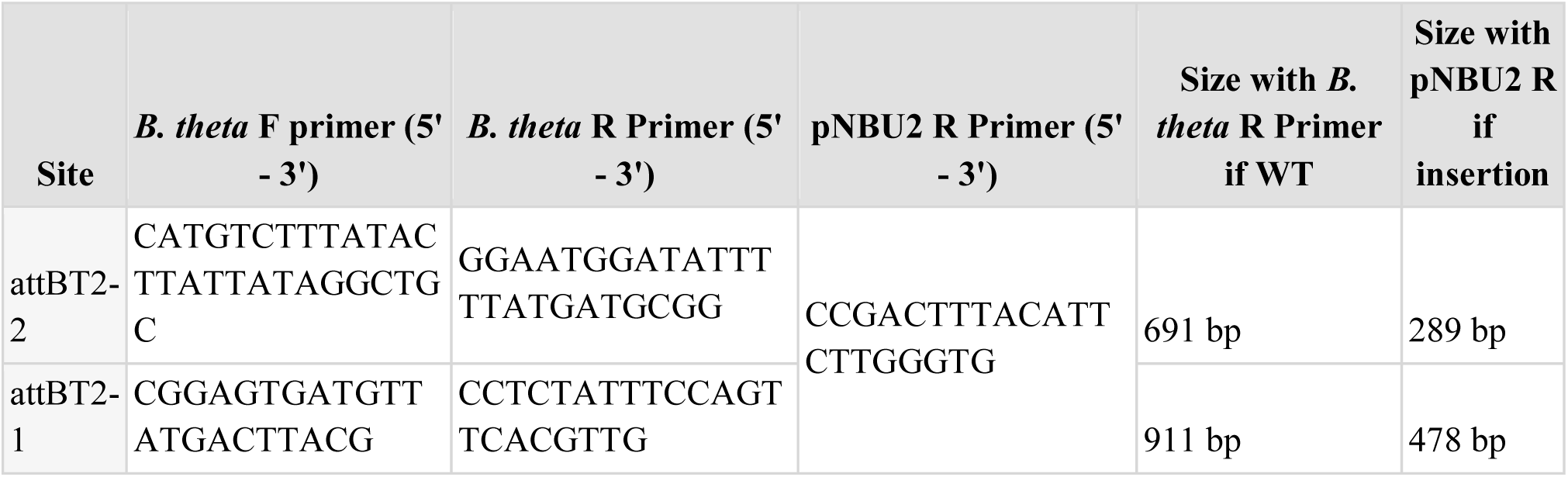
**Verification PCR for genomic insertion of the pNBU2 plasmid at the canonical attBT2 sites.** A combination of 5 primers allows identification of the pNBU2 plasmid insertion site based on amplicon length.

**Table 2.**
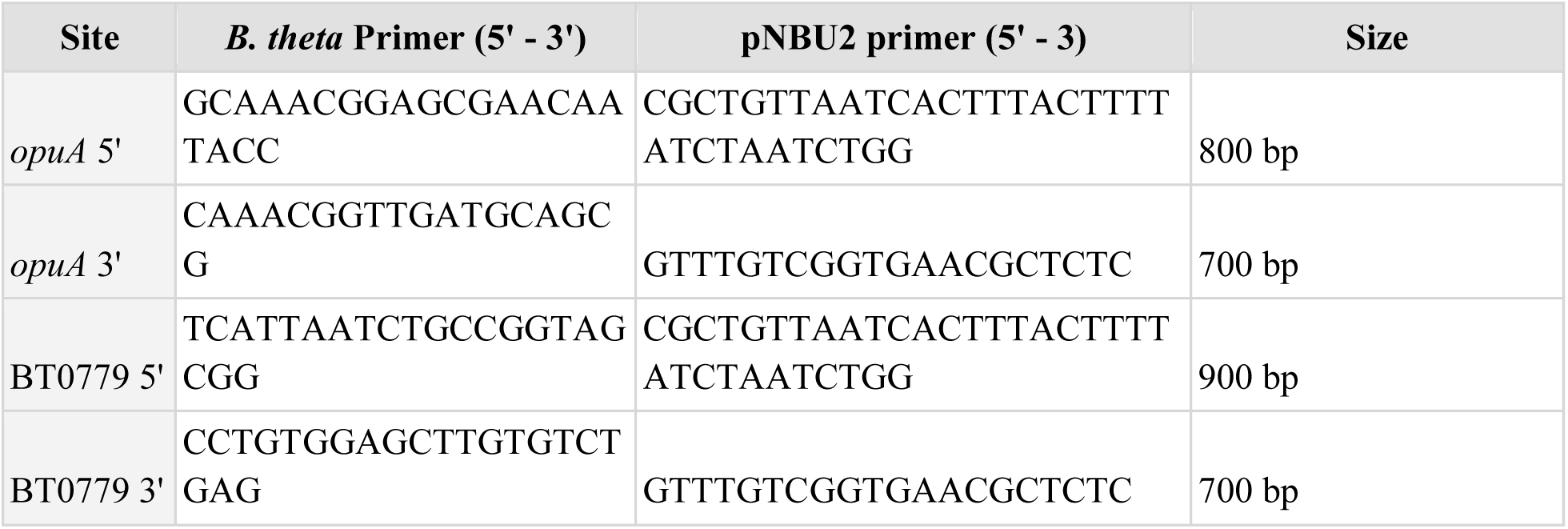
Verification PCR for genomic insertion of the biosensor plasmids at native promoter sites.

#### Genomic sequencing and analysis of engineered B. theta

*B. theta* variants of interest were struck from glycerol stocks onto selective BHIS-Erm plates and grown overnight. Single colonies were picked into fresh TYG and grown overnight. Genomic DNA was extracted from 1 mL of pelleted overnight cultures using the DNeasy PowerSoil Pro Kit (Qiagen 47014). 1 µg of purified DNA was sequenced using Oxford Nanopore sequencing (Plasmidsaurus). Genome assembly and annotation were performed by the sequencing company: (https://plasmidsaurus.com/results_interpretation#bact-assembly).

### *In vitro* fluorescence measurements

#### Promoter induction screens and orthogonality testing

For promoter library screens, glycerol stocks were thawed, diluted in fresh selective BHIS-Erm and grown 6 h at 37 °C anaerobically. Cultures were plated on BHIS-Erm plates to obtain single colonies. 189 single colonies from each library were picked and grown overnight in BHIS-Erm in 96 deep-well plates (Fisher 236600). Single colonies of a strain containing the parent P_BfP1E6_ promoter driving GFP and of the base *B. theta* strain were included in each library as controls. Overnight cultures were diluted at 1:50 into fresh BHIS containing either no inducer or small molecule inducer at the following concentrations: 100 ng/μL anhydrotetracycline hydrochloride (aTc, ThermoScientific J66688.MB); 1000 µM isopropyl β-D-1-thiogalactopyranoside (IPTG, Invitrogen 15529019); or 5 µM 2,4-diacytlphloroglucinol (DAPG, Toronto Research Chemicals D365500). Cultures were grown until they reached an OD_600_ of ∼0.3-0.7 (∼4-6 h), removed from anaerobic conditions, harvested by centrifugation for 10 min at 2500 x *g*, and resuspended twice in PBS (Fisher BP3991). After washing, cells were incubated at room temperature until they had been in aerobic conditions for a total of 2 h. 200 μL of resuspended cells were transferred to a black, clear bottom 96-well plate (Grenier Bio-One 675096), and OD_600_ and GFP levels (Ex/Em: 485/515 nm; Gain 100) were measured using a Synergy H1 plate reader (Biotek). Normalized GFP was measured as GFP/OD_600_. For orthogonality tests (Supplementary Figure 1D), strains from each library were grown as above in base media and each small molecule inducer at the levels specified above. Fluorescence was measured in a plate reader, as above. For tests comparing existing P1T_DP_ and P_LacO23_ to promoters from our libraries, strains were struck from glycerol stocks, grown overnight, and inoculated into base media. Overnight cultures were then inoculated into induced or uninduced conditions and GFP was measured, as described above.

B. theta promoter dose-response curves

Library variants of interest were struck from glycerol stocks onto selective BHIS-Erm plates and grown overnight. Single colonies were picked into fresh BHIS-Erm and grown overnight. Cultures were diluted 1:50 in 1 mL fresh BHIS supplemented with a small-molecule inducer. Small molecule inducer concentrations were as follows: 2-fold serial dilutions starting at 100 ng/mL aTc; 2-fold serial dilutions starting at 1 mM for IPTG; and 2-fold serial dilutions starting at 5μM for DAPG. The first concentration of all small-molecule inducers shown in Figure 1D,E is 0 (plain BHIS). Cultures were grown, and GFP was measured as above. Normalized GFP was averaged over three biological replicates.

### Biosensor screening and dose-response curves

Biosensor variants of interest were struck from glycerol stocks onto selective BHIS-Erm plates and grown overnight. Single colonies were picked into fresh TYG and grown overnight. Cultures were diluted 1:50 in 1 mL of fresh TYG adjusted to specified osmolalities using PEG (Bayer® MiraLAX®). To account for slower growth rates under osmotic stress, cultures at different osmolalities were diluted at staggered time intervals (approximately 10 min per 100 mOsm/kg). When appropriate, media was also supplemented with aTc at 0, 0.25, 2.5, or 25 ng/mL, IPTG at 0 or 0.5 µM, and DAPG at 0 or 0.5 µM . Cultures were grown until they reached an OD_600_ of ∼0.3-0.7, and GFP (mGreenLantern; Ex/Em: 485/515 nm; Gain: 90) and RFP (mScarlet-I3; Ex/Em: 570/600 nm; Gain 120) levels were measured. GFP and RFP were averaged over three biological replicates (three independent colonies). GFP and RFP levels were normalized to OD_600_ and to WT levels and changes in the normalized GFP:RFP ratio were measured across osmolalities.

### Biosensor osmolality induction under different growth rates

The P*_opuA_* and P*_opuA_*+20xtetO biosensors were streaked from glycerol stocks onto selective BHIS-Erm plates and incubated for two nights. Single colonies were picked into 2 mL of TYG-Erm and grown overnight. Overnight cultures were diluted 1:50 into either 1mL of fresh TYG or TYG adjusted to 510 mOsm/kg with PEG. To modulate growth rate, cultures were spiked with a gradient of SDS (Invitrogen AM9820). SDS dilutions were prepared as six consecutive 2-fold serial dilutions starting at 0.5% (w/v) and diluted 1:100 to growth media before inoculating biosensors. To track growth rate at each condition, 200 µL of each subculture was transferred into an optically clear 96-well plate (Greiner Bio-One, 655161), sealed with a clear adhesive film (Bio-Rad, 17010701), and incubated at 37 °C in a BioTek Synergy H1 plate reader. OD_600_ was measured every 5 min for 20 h with 1 min of orbital shaking between readings. In parallel, the remaining cultures were grown for ∼6 h. GFP and RFP levels were measured as described above and activation was calculated for each condition relative to baseline (340 mOsm/kg, 0% SDS) for each strain.

### *In vitro* growth curves

B. theta *growth curves* in vitro

Biosensor variants of interest were struck from glycerol stocks onto selective BHIS-Erm plates and grown overnight. Single colonies were picked into fresh TYG and grown overnight. Cultures were diluted 1:10 in 1 mL of fresh TYG and allowed to grow for 4 h to reach mid exponential phase. Cultures were then diluted again at 1:100 into 200 µL of TYG (Supplementary Figure 3G) or TYG adjusted to specified osmolalities with PEG (Bayer® MiraLAX®) (Figure 4F; Supplementary Figure 5D,E) in an optically clear 96-well plate (Greiner Bio-one 655161) and sealed with a clear seal (Bio-Rad 17010701). Cultures were then grown with continuous shaking at 37 °C in Biotek H1 synergy plate reader and OD_600_ was measured every 5 min for 20 h.

### Growth curve analysis and fitting

Individual OD_600_ curves were fitted with a Gompertz curve using a previously published in-house MATLAB script^35^. Maximum growth rate and OD_600_ were extracted using this code, and plots displaying extracted fits were created with a custom MATLAB script.

### *B. theta* transcriptomics

B. theta RNA collection from osmotic stress conditions in vitro

B. theta was struck from glycerol stocks onto BHIS plates and grown overnight. Single colonies were inoculated into pre-reduced TYG and grown overnight. Overnight cultures were then subcultured 1:50 into base TYG medium or TYG supplemented with either NaCl (Fisher S271), PEG (Bayer® MiraLAX®), lactulose (MilliporeSigma 42758450GM), or sorbitol (ACROS Organics AC132730010) to achieve osmolalities of approximately 700 and 900 mOsm/kg, which were confirmed using an Advanced Instruments Osmo1 Single-sample Micro-osmometer. Cultures were incubated until mid-exponential phase, after which samples were collected and preserved with RNAprotect (Qiagen 76506) as per manufacturer instructions. Samples were stored at −80 °C until extractions were performed.

B. theta growth in general stress conditions in vitro

B. theta was struck from glycerol stocks onto BHIS plates and grown overnight. Single colonies were inoculated into pre-reduced TYG and grown overnight. Overnight cultures were then subcultured 1:50 into the following conditions in TYG: baseline (pH 7.2); temperature stress in baseline media grown at 42 °C in a 6-well plate; acid stress adjusted to pH 5.8 with 1M HCl (Fisher SA49); oxidative stress adjusted to 100 µM H_2_O_2_ (Millipore 216763) by spiking 1M H_2_O_2_ in at a 1:100 dilution 2h and 4 h after subculturing to account for the short half-life of H_2_O_2_ in media. Cultures were incubated at either 37 °C or 42 °C until mid-exponential phase, after which samples were collected and preserved with RNAprotect as per manufacturer instructions. Samples were stored at −80 °C until extractions were done.

In vivo B. theta *transcriptomics during malabsorption*

4-5 week Swiss-Webster germ-free mice were randomly assigned to co-housed cages of the same sex, with a minimum of 3 mice per group. Mice were orally gavaged with 150 μL of overnight cultures of *B. theta* and allowed to equilibrate for 4 weeks. After equilibration, two cages (n=6) of mice were euthanized, and their cecal contents were collected to quantify bacterial abundance, measure cecal osmolality, and extract RNA. The remaining two cages (n=6) received drinking water supplemented with 15% (w/v) PEG at a final osmolality of 255-265 mOsm/kg for 6 days, as previously described^26^. On day 6 of PEG treatment, mice were euthanized, and their cecal contents were collected. Cecal samples designated for RNA extractions were flash-frozen in dry ice and stored at −80 °C until used.

### RNA extraction, library preparation and sequencing of in vitro cultures and cecal contents

*In vitro* and *in vivo* samples stored at −80 °C were thawed on ice, and their RNA was extracted using the RNeasy Mini Kit (Qiagen 74104) or RNeasy PowerMicrobiome Kit (Qiagen 26000-50), respectively, as per manufacturer instructions. RNA samples were depleted of gDNA using the TURBO DNA-free™ Kit (Invitrogen AM1907). RNA quantity and quality were verified by Qubit quantification and Agilent BioAnalyzer prior to NEBNext Ultra II RNA library preparation. Sequencing was performed on the Illumina NextSeq2000 sequencer using a P3 flow cell and 50 bp single end reads.

### Transcriptomics analysis

Transcriptomic sequencing analysis was performed using a modified script of the SAMSA2 pipeline^83^, described below. Raw reads were checked for sequencing quality using FastQC91. Following, reads were trimmed using Trimmomatic^84^ and ribosomal reads were removed from the data using SortMeRNA^85^. An indexed database was generated for *B. thetaiotaomicron* VPI-5482 using the complete reference genome (GCF_000011065.1) and raw reads were aligned to this database using DIAMOND^86^. The resulting mapped reads were analyzed on DESeq2^30^ to perform differential expression analysis. Differential gene expression was performed using *B. theta* samples grown in base TYG or samples from untreated mice as the reference condition.

### Mouse Experiments and *in vivo B. theta* fluorescence measurements

#### 0%, 10%, and 15% PEG dosage mouse experiments (Figures 4A, 5A)

3–5-week-old Swiss-Webster germ-free mice were randomly assigned to co-housed cages of the same sex. Mice were orally gavaged with 150 μL of overnight cultures of *B. theta* biosensor strains P*_opuA_*+20xTetO or P_BT0779_+160xTetO, or the No Promoter control strain. Three cages were inoculated per strain (n = 3-5 mice per cage). After a 2-week equilibration period, mice were started on either 0%, 10%, or 15% (w/v) PEG in drinking water, at final osmolalities of 0, 104, and 216 mOsm/kg, respectively, for 6 days. Upon completing 6 days, mice were euthanized, and their cecal contents were collected and used for osmolality measurements and flow cytometry analysis.

*0-7.5% PEG dosage mouse experiment (Figure 5D)*

11–14-week-old Swiss-Webster germ-free mice were randomly assigned to co-housed cages of the same sex, with 3-5 mice per cage. All mice were orally gavaged with 150 μL of overnight cultures of P_BT0779_+160xTetO. After a 2-week equilibration period, mice were started on either 0%, 1%, 2.5%, 5%, or 7.5% (w/v) PEG in drinking water at final osmolalities of 0, 5, 14, 35, and 66 mOsm/kg, respectively, for 3 days. On Day 3, fecal pellets were collected from each mouse and used for flow cytometry and Bristol Stool Scoring^47^. The same fecal pellets were used to measure fecal water content as previously described. Briefly, microfuge tubes were weighed using an analytical balance before fecal pellets were added^46^. The combined weight was recorded, and the samples were dried overnight at 60 °C with tube lids open. The final tube weight was then recorded, and fecal water content was calculated as: (fecal weight before drying − fecal weight after drying)/feces weight before drying × 100. On the same day as fecal sample collection, mice were euthanized, and their cecal contents were collected and used for osmolality measurements and flow cytometry analysis.

### 15% and 20% PEG dosage mouse experiments (Figure 4G)

14-week-old female Swiss-Webster germ-free mice were co-housed in the same cage. Mice (n=5) were orally gavaged with 150 μL of overnight cultures of *B. theta* biosensor strains P*_opuA_*+20xTetO^HR^. After a 1-week equilibration period, mice were started on 20% (w/v) PEG in drinking water, at final osmolality of 379 mOsm/kg, for 6 days. Fecal samples were collected on days 0, 2, 4, and 6 of the treatment and processed for flow cytometry. After 6 days of treatment, mice were euthanized, and their cecal contents were collected and used for osmolality measurements and flow cytometry analysis.

### B. theta abundance measurements from mono-colonized mice

Bacterial colony-forming units (CFU) were quantified by sampling cecal contents with 1 µL loops and resuspending in 200 μL of PBS. Ten-fold serial dilutions of the resuspended samples were performed in PBS, and 5 μL of each dilution was spot-plated in BHIS agar. Plates were incubated until single colonies were countable (∼24 h).

### Biosensor abundance measurements in a complex microbiota

Bacterial CFUs were quantified by sampling fecal pellets with 1 µL loops and resuspending in 100 μL of PBS. Ten-fold serial dilutions of the resuspended samples were performed in PBS, and 5 or 25 μL of each dilution was spot-plated on BHIS agar supplemented with Erm and Gent. Plates were incubated anaerobically until single colonies were countable (∼24 h), then stored aerobically at 4 °C prior to quantification. Because other microbiota members displayed resistance to the selection antibiotics, biosensor density was determined specifically from GFP^+^ or RFP^+^ colonies using a Zeiss Stemi SV11 fluorescent dissecting microscope.

### Cecal osmolality measurements

Cecal contents collected when mice were euthanized were centrifuged at 16,000 x *g* for 20 min at 4 °C immediately after collection. The resulting cecal supernatant was used to measure cecal osmolality using an Advanced Instruments Model 3220 osmometer.

### Isolation of live bacteria from fecal and cecal contents of mono-colonized mice

Live bacteria isolation protocol was adapted from a previous protocol^87^. One fourth of fecal pellets (∼10-20 mg), or ∼20-50 mg of cecal contents, were placed in 2 mL microfuge tubes 1-2 h after collection. One stainless steel bead (5.0 mm) was added to each tube together with 1 mL of PBS. The samples were thoroughly homogenized using TissueLyser II (Qiagen) for 10 min at 20 Hz. The homogenized samples were then subjected to three iterations of the following steps: vortexed for 15 s; centrifuged at 95 x *g* for 30 s at room temperature; 750 µL of supernatant transferred into a fresh 2 mL microcentrifuge tube pre-filled with 750 µL PBS. After the final iteration, tubes were centrifuged at 4,000 x *g* for 5 min at room temperature, the supernatant was discarded, and pellets were resuspended in 1 ml of PBS. Resuspended samples were then diluted 1:10 in PBS in preparation for flow cytometry. Isolations were performed aerobically to allow for fluorescent proteins to mature.

### Isolation of live bacteria from fecal contents of conventional mice

One fecal pellet (∼30-50 mg) was placed in 2 mL microfuge tubes 1-2 h after collection. One stainless steel bead (5.0 mm) was added to each tube together with 1 mL of PBS. The samples were thoroughly homogenized using TissueLyser II (Qiagen) for 10 min at 20 Hz. The homogenized samples were then subjected to 4 iterations of the following steps: vortexed for 15 s; centrifuged at 200 x *g* for 5 min at room temperature; supernatant transferred into a fresh 2 mL microcentrifuge tube. After the final iteration, tubes were subjected to two iterations: centrifugation at 5,000 x *g* for 5 min at room temperature, supernatant discarded, and pellets resuspended in 1 ml of PBS. Finally, tubes were centrifuged one last time at 200 x *g* for 10 min, and the supernatant was transferred and diluted 1:2 in PBS in preparation for flow cytometry. Isolations were performed aerobically to allow for fluorescent proteins to mature.

### Flow cytometry protocol and analysis for mono-colonization mouse experiments

Isolated *B. theta* biosensors from fresh fecal or cecal samples were analyzed on a CytoFLEX LX flow cytometer using the following excitation laser and emission filter combinations for: mGL (Ex/Em: 488/525 nm) and mScarlet-I3 (Ex/Em: 488/610 nm). Samples were diluted 1:10 or 1:100 in PBS for optimal detection of fluorescence. Single-fluorophore *B. theta* controls (GFP^+^RFP^-^ and GFP^-^RFP^+^) were grown in TYG, and 1 mL samples were collected at mid-exponential phase (OD_600_ ∼0.5), pelleted at 5,000 x *g* for 5 min, resuspended in 1 mL of PBS, and mixed with a germ-free fecal sample processed as explained above. These controls were used to establish acquisition gating parameters that distinguish bacterial fluorescence from background fecal fluorescence. An acquisition gate was set at an mScarlet threshold >4×10 arbitrary units (AU), and acquisition was terminated once 10,000 events within this gate were recorded. All samples were recorded at a low-speed flow rate (10 µL/min). FCS files were imported into FlowJo (v10.10.0), where mScarlet fluorescence was overlaid on all events in the forward and side scatter channels and a gate (“Cells”) was set around the area with the highest mScarlet fluorescence (Supplementary Figure 5A). Events within this “Cells” gate were then gated based on the events that had mScarlet^+^ fluorescence (>2×10^3^ AU), separating our biosensor events from background fluorescence (Supplementary Figure 5B). All events within the mScarlet^+^ gate for each sample were exported, and their fluorescence output values were analyzed and visualized with R using tidyverse (v2.0.0) and ggridges (v0.5.6) packages.

### Flow cytometry protocol and analysis for conventional mouse experiments

Bacteria isolated from fecal pellets were analyzed on a 5-laser Aurora spectral analyzer (Cytek Biosciences) using the standard manufacturer optical configuration. To enable autofluorescence spectral unmixing, a background fecal sample from non-colonized mice was processed using the same isolation procedure as described above and acquired in parallel. Autofluorescence signatures were defined for this background population using the autofluorescence explorer in the Cytek software. Single-fluorophore *B. theta* controls (GFP^+^RFP^-^ and GFP^-^RFP^+^) and the P_BT0779_+160xTetO biosensor were cultured in TYG medium to mid-exponential phase, pelleted, resuspended in PBS, mixed with the background fecal sample, and acquired. Based on these controls, an acquisition gate was set at an RFP threshold >7×10^3^ A.U. to reduce auto-fluorescent fecal background collected. For each experimental sample, acquisition was stopped once either 50,000 gated events or 1.5 mL of sample had been collected. All samples were recorded at a low flow rate (10 µL/min). FCS files were imported into FlowJo, where biosensor populations were gated on RFP^+^ and autofluorescence channels (Supplementary Figure 6G). Events within this gate were exported for each mouse, and fluorescence outputs were analyzed and visualized with R using tidyverse and ggridges packages.

### Biosensor Computational Modelling

#### Model construction

The following ODE-based mathematical models representing 1) P_TetR56_ induction with aTc (Equations 1-2) and 2) native promoter biosensor induction in response to a natural stimulus (Equations 3-6) were constructed in MATLAB (MathWorks) and solved using the ode23.m function:

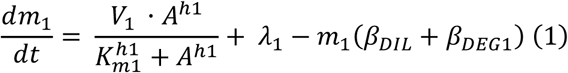

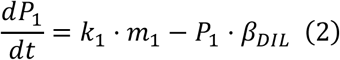

where:

𝑚_1_ is the number of GFP mRNAs

𝑃_1_ is the number of GFP proteins

𝑉_1_ is the maximum transcription rate from P_TetR56_

𝐴 is the concentration of aTc

ℎ1 is the Hill coefficient of the P_TetR56_ promoter

𝐾_𝑚1_ is the repression threshold of the P_TetR56_ promoter, at which expression is half-maximal

## 𝜆_1_ is the leaky expression rate constant for the P_TetR56_ promoter

## 𝛽_𝐷𝐼𝐿_ is the rate constant of dilution due to cell growth

## 𝛽_𝐷𝐸𝐺1_ is the degradation rate of GFP mRNA

𝑘_1_ is the translation rate constant of GFP protein from GFP mRNA

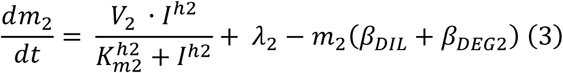

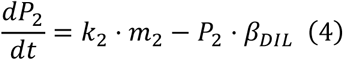

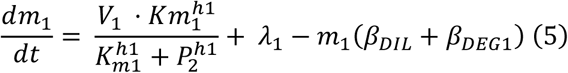

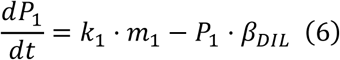

where:

𝑚_1_ is the number of GFP mRNAs

𝑃_1_ is the number of GFP proteins

𝑚_2_ is the number of TetR mRNAs

𝑃_2_ is the number of TetR proteins

𝑉_1_ is the maximum transcription rate from P_TetR56_

ℎ1 is the Hill coefficient of the P_TetR56_ promoter

𝐾_𝑚1_ is the repression threshold of the P_TetR56_ promoter, at which expression is half-maximal

𝜆_1_ is the leaky expression rate constant for the P_TetR56_ promoter

𝛽_𝐷𝐼𝐿_ is the rate constant of dilution due to cell growth

𝛽_𝐷𝐸𝐺1_ is the degradation rate of GFP mRNA

𝑘_1_ is the translation rate constant of GFP protein from GFP mRNA

𝑉_2_ is the maximum transcription rate from the native *B. theta*

𝐼 is the concentration of inducer of the native promoter

ℎ2 is the Hill coefficient of the native promoter

𝐾𝑚_2_ is the induction threshold of the native promoter

𝜆_2_ is the leaky expression rate constant for the native promoter

𝛽_𝐷𝐸𝐺2_ is the degradation rate of TetR mRNA

𝑘_2_ is the translation rate constant of TetR protein from TetR mRNA

Hill dynamics were assumed for repressible transcription from the P_TetR56_ and inducible transcription from native *B. theta* promoters. Sponging of TetR follows Hill ligand binding dynamics, with the degree of sponging varying with TetR concentration and number of total binding sites included (*S*). As TetR dimerizes before binding operator sites, the TetR sponging term is multiplied by 2. We assumed that no active degradation of TetR or GFP occurs without degradation tags added.

### Parameter fitting

The Covariance Matrix Adaptation-Evolution Strategy (CMA-ES) algorithm^88^ was employed to fit parameter values for the above models that reflect system behaviour for P_TetR56_ and both P*_opuA_* and P_BT0779_ biosensors. Here, CMA-ES works by stochastically exploring the spaces around a given set of estimated parameter values. At each set of parameter values, a cost is calculated as a function of distance between model results with the current parameter set and a user-provided experimental dataset. Over a defined number of iterations, CMA-ES tries to minimize the calculated cost, homing in on a parameter set that represents the closest fit to the experimental data provided.

In this case, initial parameter values were estimated based on biologically relevant transcription and translation rates from *E. coli*^40,89,90^, and CMA-ES calculated cost based on the distance between predicted steady-state GFP levels at a given parameter set and experimental GFP measurements from *B. theta* dose response curves. Specifically, CMA-ES (cmaes.m^91^) was first run on Model 1 using the aTc dose response curve (Figure 1D) to fit the following parameters: 𝜆_1_, 𝑉𝑚𝑎𝑥_1_, 𝐾𝑚_1_, 𝐵𝑒𝑡𝑎_𝑑𝑒𝑔_, 𝑘_1_. Using these estimates as initial input parameters for Model 2, CMA-ES was then run on Model 2 without sponges (*S* = 0), using osmolality induction curves from P*_opuA_* biosensors as input data to fit the remaining parameters. In both cases CMA-ES was run 25 times for 500 iterations each. The parameter set with the lowest cost from all runs combined was saved as the optimized parameter set to use for modelling the system with sponging. Initial estimated and optimized parameter values for both biosensors are listed in Supplementary Table 5. Using the parameter sets estimated by CMA-ES for P*_opuA_*, Model 2 was then solved over a range of induction levels (*I*) and repressor sponging (*S*). Steady-state GFP values from each solution were used to calculate fold change in GFP across conditions.

### Degradation and growth rate simulations

To include constative degradation of GFP to the model, an extra term was included in Equation 6 to form the following equation defining GFP protein dynamics:

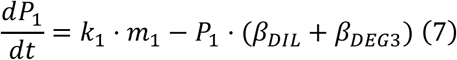

Where 𝛽_𝐷𝐸𝐺3_ is the degradation rate constant of GFP. When biosensor dynamics were assessed across degradation and growth rates, values for 𝛽_𝐷𝐸𝐺3_ and 𝛽_𝐷𝐼𝐿_ were altered iteratively, and biosensor activation, baseline steady-state GFP levels, and the time to half-maximal GFP levels were assessed at each iteration.

### Statistical Analyses

Statistical analyses performed on given data sets are indicated in the corresponding figure legends. For all figures: * *padj* < 0.05; ** *padj* < 0.01; *** *padj* < 0.001.

## Acknowledgments

The authors acknowledge that the land we performed this research on is the traditional, ancestral, and unceded territory of the xwməθkwəy̓əm (Musqueam) Nation. The land it is situated on has always been a place of learning for the Musqueam people, who for millennia have passed on in their culture, history, and traditions from one generation to the next on this site. We encourage others to learn more about the native lands in which they live and work at https://native-land.ca/. The authors acknowledge support from the following: CIFAR / Humans and the Microbiome (FL-001253 Appt 3362, to C.T.); Michael Smith Foundation for Health Research Scholar Award (18239, to C.T.); Canada Foundation for Innovation / Infrastructure Operating Fund (38277); Canada Tier 2 Research Chair, Quantitative Microbiota Biology for Health Applications (CRC-2022-00036, to C.T.); Weston Family Foundation / Weston Family Microbiome Initiative (F21-05234, to C.T., L.P.T.); NSERC Discovery grant (RGPIN-2019-04591, to C.T.); CIFAR Azrieli Global Scholars (GS20-011, to C.T.); NSERC-CREATE SynBioApps Fellowship (to G.M.); NSERC Canada Graduate Scholarships (to G.M.). This work received support from the Centre of Disease Modelling, especially Natalia Caranza, and from ubcFLOW. This research was supported in part through computational resources and services provided by Advanced Research Computing (ARC) at the University of British Columbia. The authors also thank Michael Hunter for critically reading this manuscript and providing feedback, Tatiana Lau for her help with FACS, as well as Claire Sie and Apsara Srinivas for their help with animal work.

## CRediT authorship contribution statement

GM, JCB: Conceptualization, Methodology, Software, Validation, Formal analysis, Investigation, Writing

- Original Draft, Writing - Review & Editing, Visualization

JH: Conceptualization, Methodology, Investigation, Formal analysis AH: Investigation

LPT: Conceptualization, Methodology, Resources, Writing - Review & Editing, Supervision, Project administration, Funding acquisition

CT: Conceptualization, Methodology, Resources, Writing - Review & Editing, Supervision, Project administration, Funding acquisition

---

**Supplementary Figure 1.**
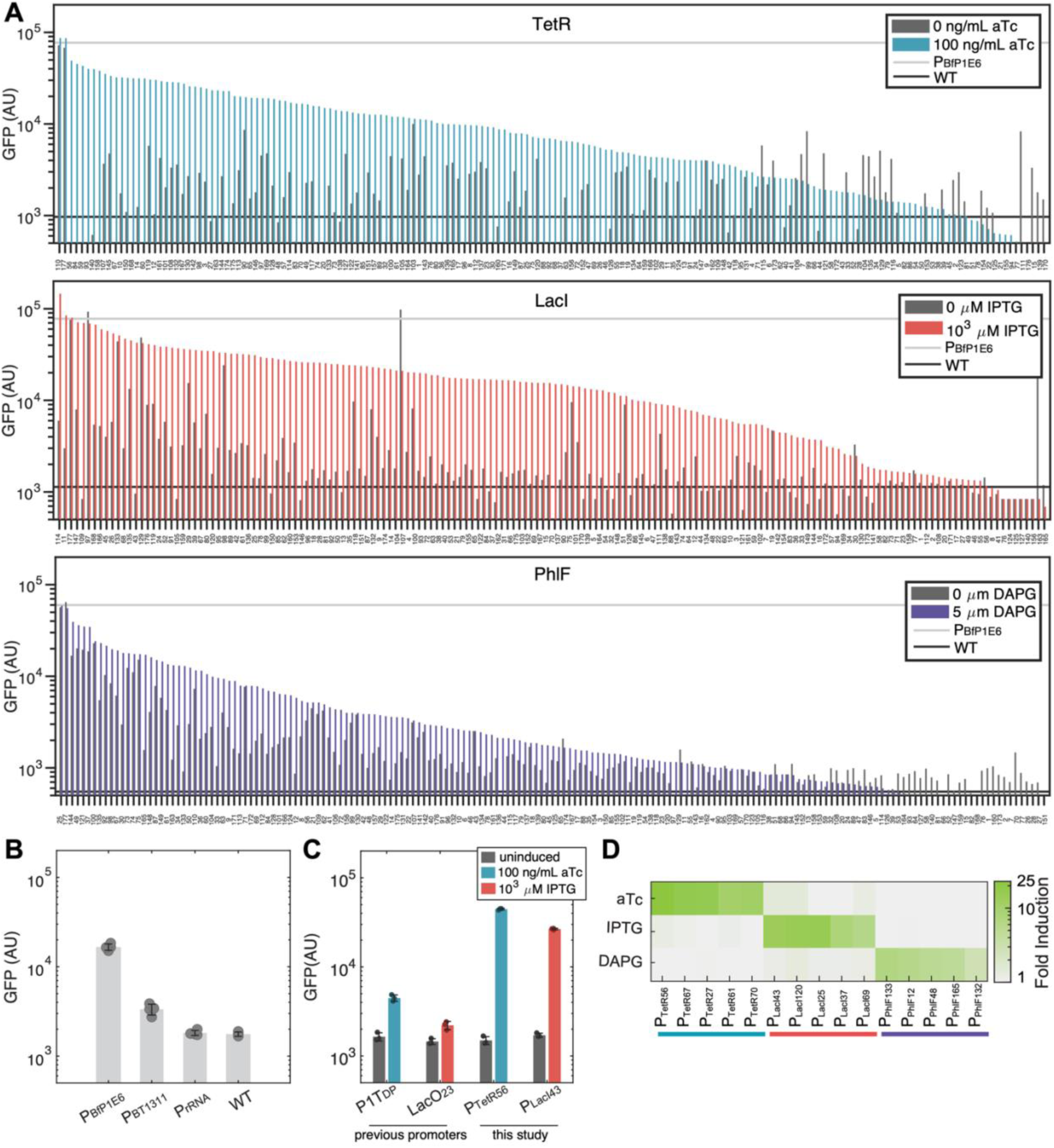
**Engineered phage promoters enable strong, repressible, and orthogonal control of fluorescence expression in *B. theta*.** A) Activity of TetR, LacI, and PhlF promoter variants measured by GFP expression in uninduced (base media) and induced conditions (100 ng/mL aTc, 10^3^ μM IPTG, or 5 μM DAPG, respectively). Light and dark horizontal lines represent the parent phage promoter and wild type *B. theta*, respectively, in media with an inducer present. B) GFP levels driven by strong native *B. theta* promoters compared to the parent P_BfP1E6_ phage promoter used in our promoter libraries. C) GFP levels driven by previously developed TetR- and LacI-regulated *B. theta* promoters (P1T_DP_^8^ and P_LacO23_^7^) compared to the P_TetR56_ and P_LacI43_ promoters from this study in their corresponding maximal induction conditions (100 ng/mL aTc or 10^3^ μM IPTG, respectively). D) Orthogonality matrix of strong inducible promoters from each of the three libraries. Strains were incubated in 100 ng/mL aTc, 10^3^ μM IPTG, or 5 μM DAPG, and the fold-change GFP was calculated compared to uninduced conditions. B,C: Error bars represent the standard deviation from n = 3 biological replicates (unique colonies) of each promoter. D: Heatmap values represent the mean fold-change GFP values from 3 biological replicates.

**Supplementary Figure 2.**
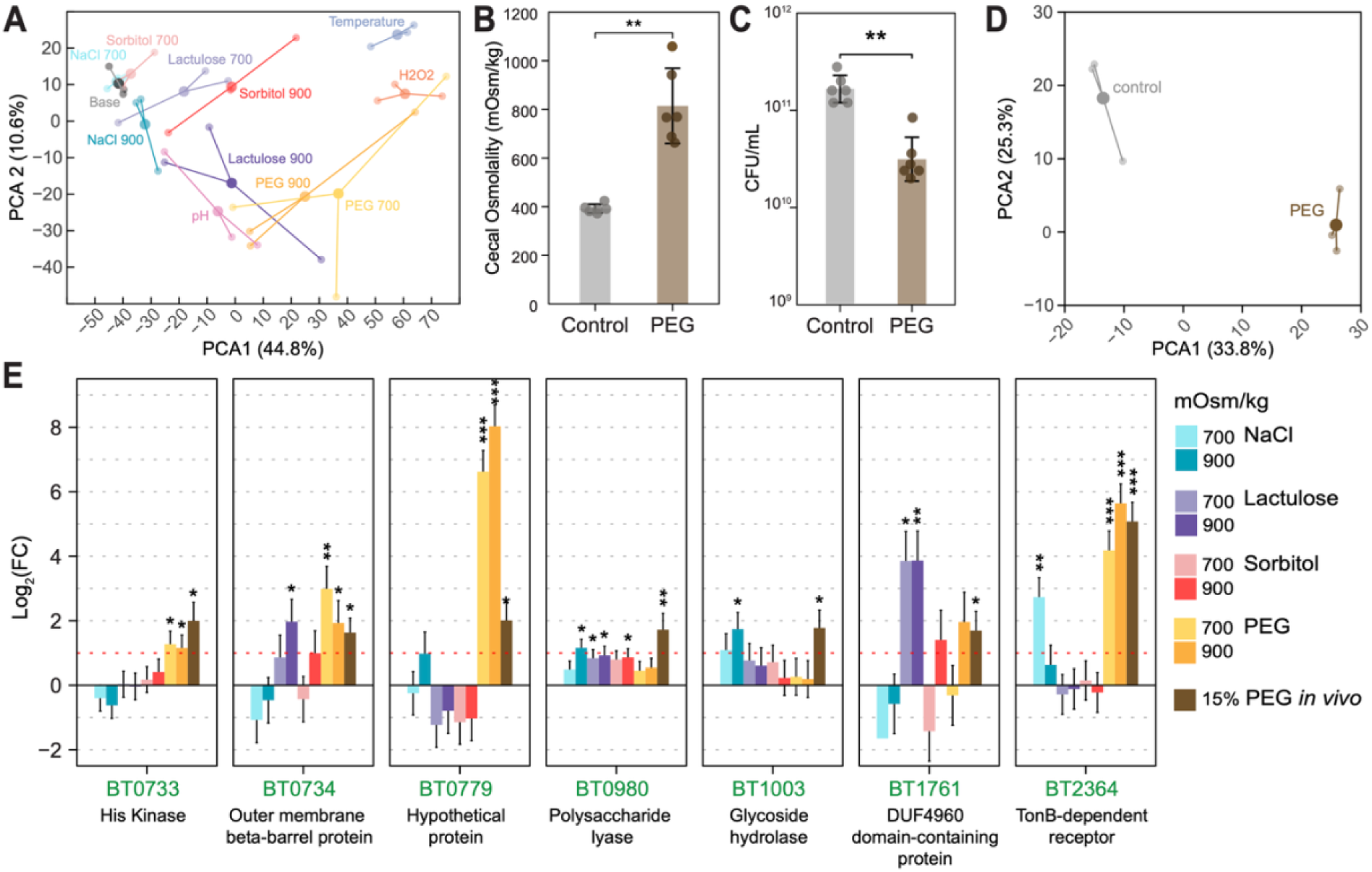
**The *B. theta* transcriptional landscape is altered by osmolality *in vitro* and *in vivo*.** A) Principal component analysis (PCA) of the *B. theta* transcriptome after treatment with NaCl, lactulose, PEG, or sorbitol *in vitro*. Small circles indicate the coordinates of each biological replicate, large circles indicate the mean position of three biological replicates, and lines mark the Euclidean distance from each replicate to the mean (shown in Figure 2A). B) Cecal osmolality of mice treated with 15% PEG for 6 days, or untreated controls. C) Bacterial abundance counts from mice treated with 15% PEG for 6 days, or untreated controls. D) PCA of the *B. theta* transcriptome from mono-colonized mice treated with 15% PEG for 6 days and untreated controls. E) Differential expression analysis of the 7 genes identified that are specifically upregulated in osmotic stress conditions *in vitro* and *in vivo* (highlighted by the dotted line in Figure 2F). Log_2_(FC) of the genes in each *in vitro* osmotic stress condition and following 6 days of treatment with 15% PEG are shown. B,C: Error bars indicate the standard deviation of *n* = 3 biological replicates. Pairwise ANOVA with Tukey’s multiple correction (* *p* < 0.05; ** *p* < 0.01; *** *p* < 0.001) E: Error bars indicate the standard error of *n* = 3 biological replicates. Statistics calculated in DESeq2 pipeline^30^ (* *padj* < 0.05; ** *padj* < 0.01; *** *padj* < 0.001)

**Supplementary Figure 3.**
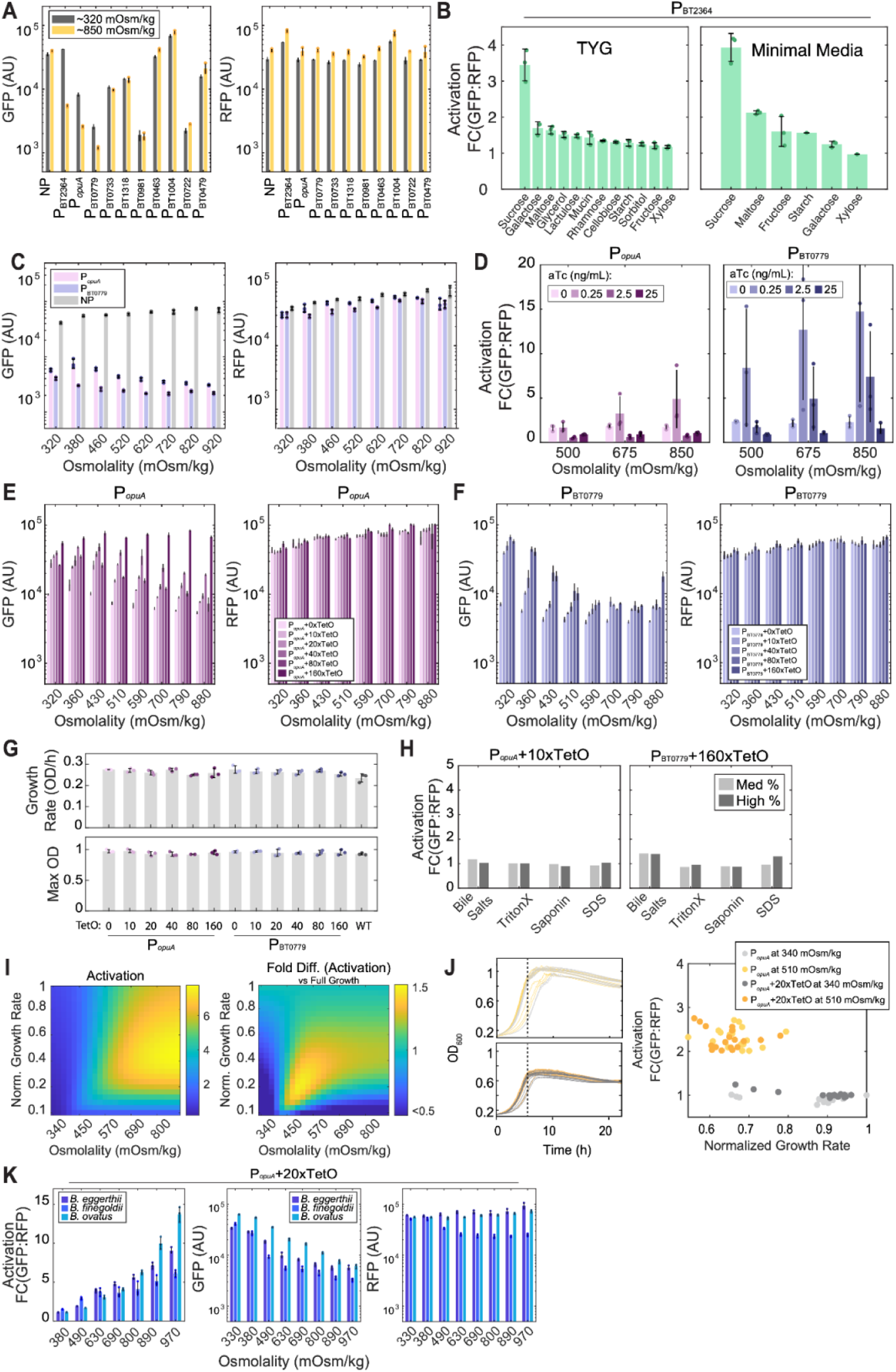
**Biosensor response is enhanced by repressor sequestration, specific to osmolality, relatively robust to growth rate, and portable across *Bacteroides*.** A) GFP and RFP levels for each biosensor shown in Figure 3B in media adjusted to 320 mOsm/kg or 850 mOsm/kg with PEG. B) Activation of the P_BT2364_ biosensor when grown in (*left*) TYG or (*right*) minimal media with indicated carbohydrates supplemented at 0.5%. In minimal media conditions, no other carbon source was added to support growth. Activation is calculated relative to cultures grown in 0.5% glucose as a carbon source. C) P*_opuA_*and P_BT0779_ biosensor GFP and RFP levels from experiment shown in Figure 3C in media adjusted to increasing osmolalities with PEG. D) Biosensor response to osmolality with chemical sequestration of TetR via aTc supplementation. P*_opuA_* and P_BT0779_ biosensors were grown in media adjusted to specified osmolalities with PEG, and supplemented with 0, 0.25, 2.5, or 25 ng/mL aTc, and GFP:RFP ratios were compared to baseline media (∼320 mOsm/kg). E) P*_opuA_* biosensor GFP and RFP levels from the TetR sponging experiment in PEG shown in Figure 3G. F) P_BT0779_ biosensor GFP and RFP levels from the TetR sponging experiment in PEG shown in Figure 3G. G) Maximum growth rate and OD from growth curves of P*_opuA_* and P_BT0779_ biosensor sponge variants. Cultures were grown in TYG at 330 mOsm/kg for 24 h. The number of sponges for each variant is indicated. H) P*_opuA_*+10xTetO response measured in media supplemented with membrane-damaging agents at the following concentrations (Med % or High %): Bile salts (0.25% or 0.5% w/v), Triton-X (0.0031% or 0.0063% v/v), saponin (0.025% or 0.05% w/v), SDS (0.0016% or 0.0031% v/v). For Triton-X and saponin, concentrations higher than those listed did not allow cell growth. I) Simulations of the P*_opuA_* biosensor across induction levels and growth rates predicts that growth rate can increase or decrease biosensor output at a given induction level. The maximum difference in activation due to growth rate is predicted at ∼520 mOsm/kg, under substantially slowed growth. J) *In vitro* experiments confirm some variation across growth rate when biosensors were grown in slightly induced conditions (510 mOsm/kg) and growth rate was slowed independent of osmolality using SDS. Here, (*left*) growth rate was measured from growth curves in each condition, and (*right*) biosensor activation was measured after 6h. A maximum ∼1.4-fold variation in activation was observed in induced conditions. K) Biosensor activation, GFP, and RFP levels for the P*_opuA_+*20xTetO biosensor hosted in indicated *Bacteroides* strains. TYG growth media was adjusted to indicated osmolality with PEG. A,B,C,D,E,F,G,K: Error bars represent the standard deviation from *n* = 3 biological replicates (unique colonies) of biosensors in each condition. Where shown, individual points represent individual biological replicates. H: Bar height indicates the value from a single biological replicate.

**Supplementary Figure 4.**
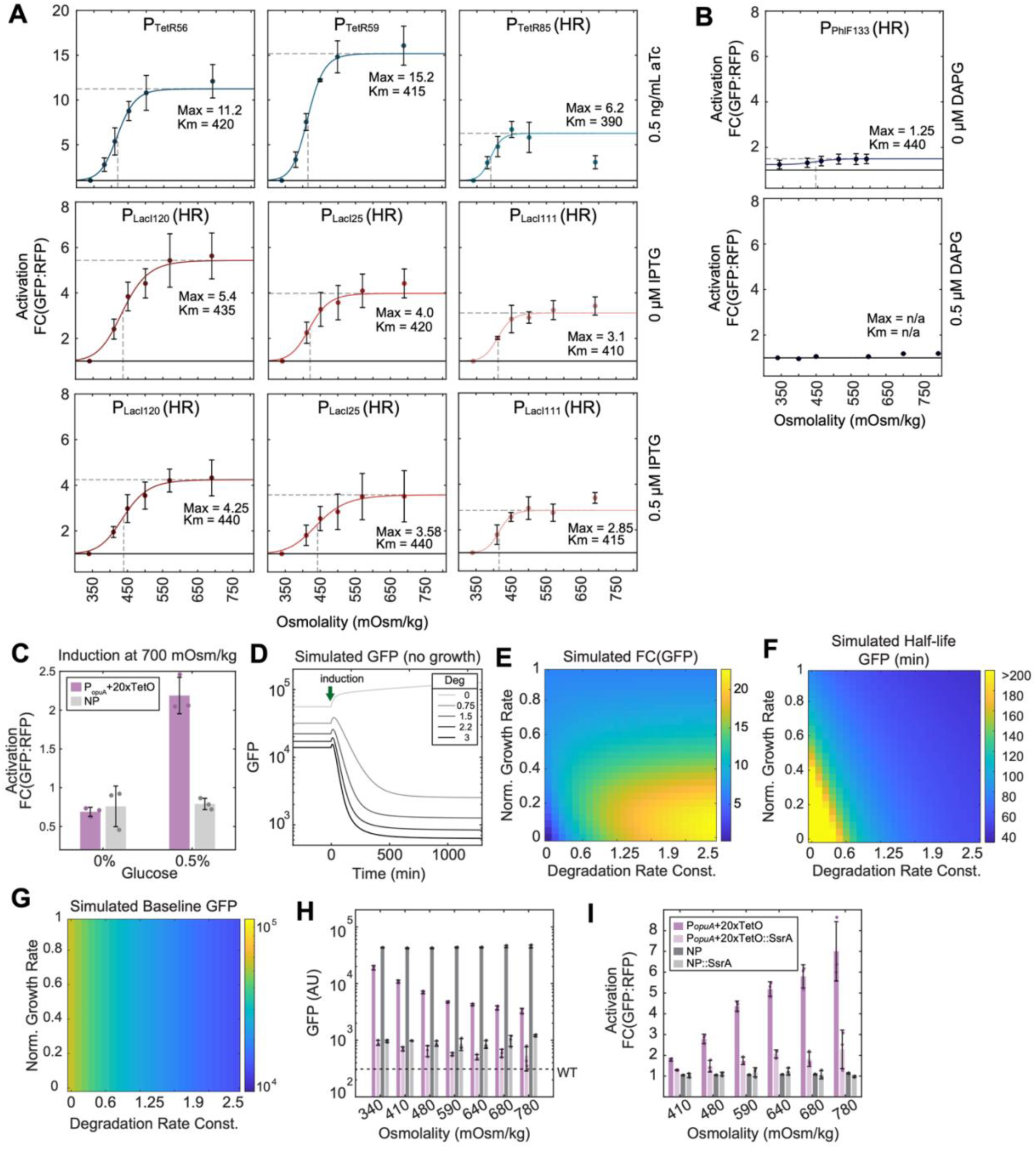
**Repressible promoter selection and targeted GFP degradation can tune biosensor activation and sensitivity, and is predicted to improve response dynamics.** A-B) Induction curves for P_BT0779_ biosensor variants constructed using diverse inducible promoters regulated by (A) P_TetR_, P_LacI_, or (B) P_PhlF_ repressors. P_TetR_ biosensor variants were grown with 0.5 ng/mL aTc to mimic optimal repressor sequestration that maximized dynamic range (Supplementary Figure 3D). P_LacI_ and P_PhlF_ biosensor variants were grown with and without 0.5 µM IPTG or DAPG, respectively. P_LacI_ variant responses were decreased with repressor sequestration, while P_PhlF_ biosensors showed no response with repressor sequestration. Osmolality of all growth media was adjusted with PEG. Where shown, solid lines represent the fit curve for the first 5 osmolalities, fit to a Hill equation with fit.m in MATLAB. For each biosensor variant, dashed horizontal lines represent the fit maximal activation, and dashed vertical lines represent the fit *K_m_*. HR indicates variants that are conjugated by homologous recombination at the native BT0779 locus. C) Induction of the P*_opuA_*+20xTetO biosensor in minimal media at 700 mOsm/kg with and without a carbon source. Growth is required for biosensor response. D) Simulations of the biosensor response, including targeted degradation of GFP. Even without growth, exponential decay of GFP is predicted. E) Predicted fold decrease of GFP at maximum induction (1000 mOsm/kg) across a range of growth rate and degradation rate constants. F-G) Phase space showing simulated (F) half-life of GFP signal and (G) baseline GFP levels across a range of growth rate and degradation rate constants. Simulations are run at 1000 mOsm/kg induction. Overall, increasing degradation leads to decreased GFP half-life, but also to lower baseline GFP signal. H) GFP levels of the P*_opuA_*+20xTetO biosensor and NP control, as well as SsrA-tagged variants of each construct, at increasing osmolalities. Fusion to the SsrA tag leads to strong degradation of GFP in both constructs. Osmolality of media was adjusted with PEG. I) Activation levels of the P*_opuA_*+20xTetO biosensor and NP control and their respective SsrA-tagged variants in increasing osmolalities, from results shown in H. The P*_opuA_*+20xTetO::SsrA biosensor showed slight activation compared to the NP::SsrA control, but dynamic range was decreased compared to the untagged control. A,B,C,H,I: Individual points represent 3 biological replicates (individual colonies). Error bars show standard deviation between replicates.

**Supplementary Figure 5.**
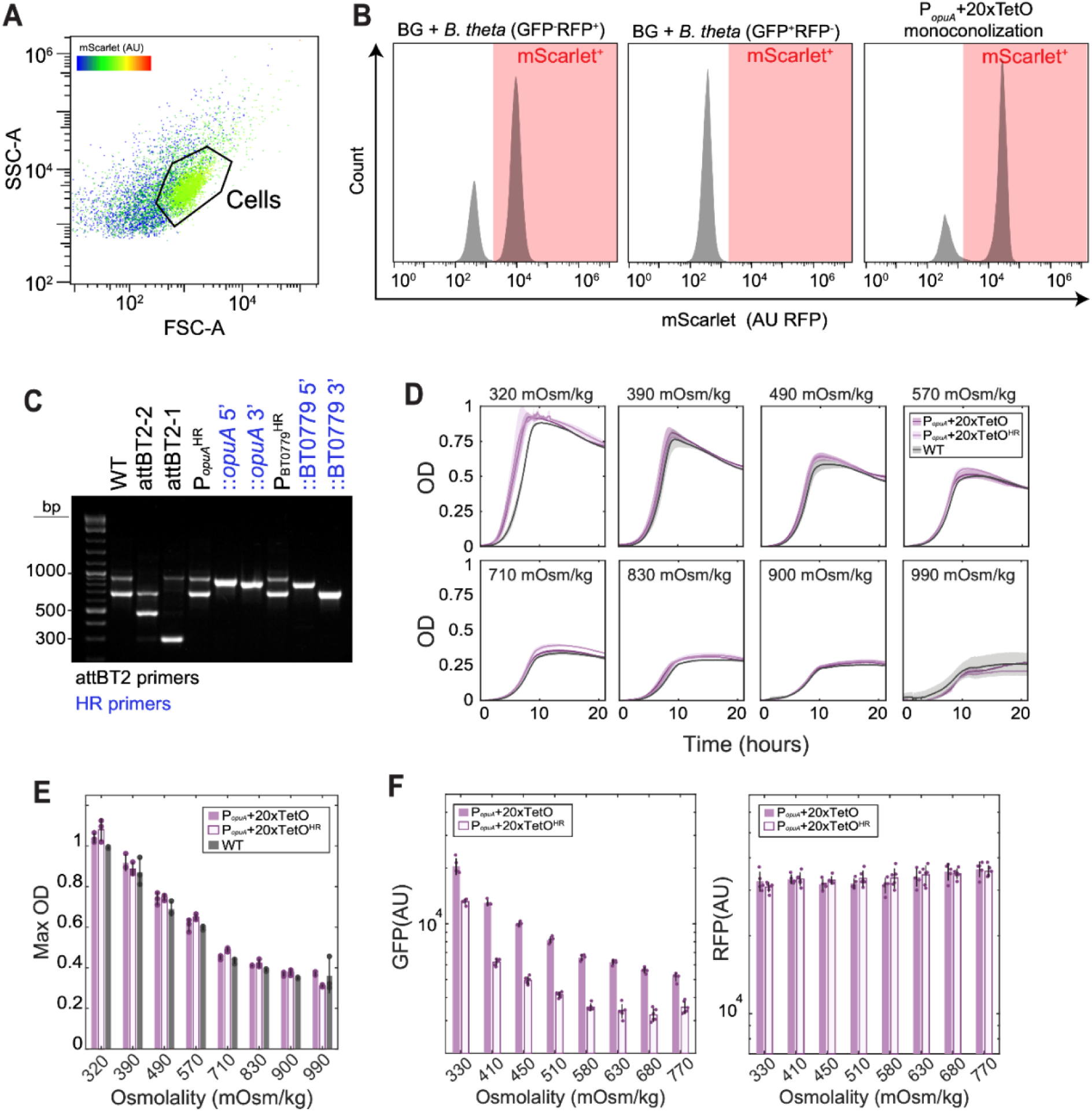
**Integration of transcriptional reporter circuits can occur via homologous recombination without affecting fitness.** A-B) Flow cytometry gating for biosensors from fecal or cecal samples of mono-colonized mice. A) Biosensor cell populations were gated from fecal preparations based on mScarlet (RFP) fluorescence density. B) Gating strategy for identifying biosensors from fecal and cecal autofluorescence in mono-colonized mice. Germ free fecal samples (background; BG) were spiked with (*left*) GFP^-^ RFP^+^ or (*middle*) GFP^+^ RFP^-^ strains grown *in vitro* to define an appropriate threshold for identifying cells. This threshold was then applied to (*right*) an experimental sample from a representative mouse mono-colonized with P*_opuA_*+20xTetO biosensor to verify gating. C) Agarose gel electrophoresis showing (*black labels*) PCR products from the three possible insertion sites or wild type *B. theta* using our insertion verification primers, and (*blue labels*) PCR products from primer pairs verifying insertion of the P*_opuA_*+20xTetO^HR^ or P_BT0779_+160xTetO^HR^ biosensor circuits via homologous recombination at their respective native promoter sites. D) OD_600_ measurements from growth curves for P*_opuA_*+20xTetO and P*_opuA_*+20xTetO^HR^ biosensors in media adjusted to increasing osmolalities with PEG. E) Max OD_600_ values from growth curves shown in D. F) GFP and RFP values normalized to OD_600_ from dose response curves in Figure 4D. D: Solid lines and shaded error bars represent the mean and standard deviation from *n* = 3 biological replicates (unique colonies) of biosensors in each condition. E,F: Error bars represent the standard deviation from *n* = 3 biological replicates (unique colonies) of biosensors in each condition.

**Supplementary Figure 6.**
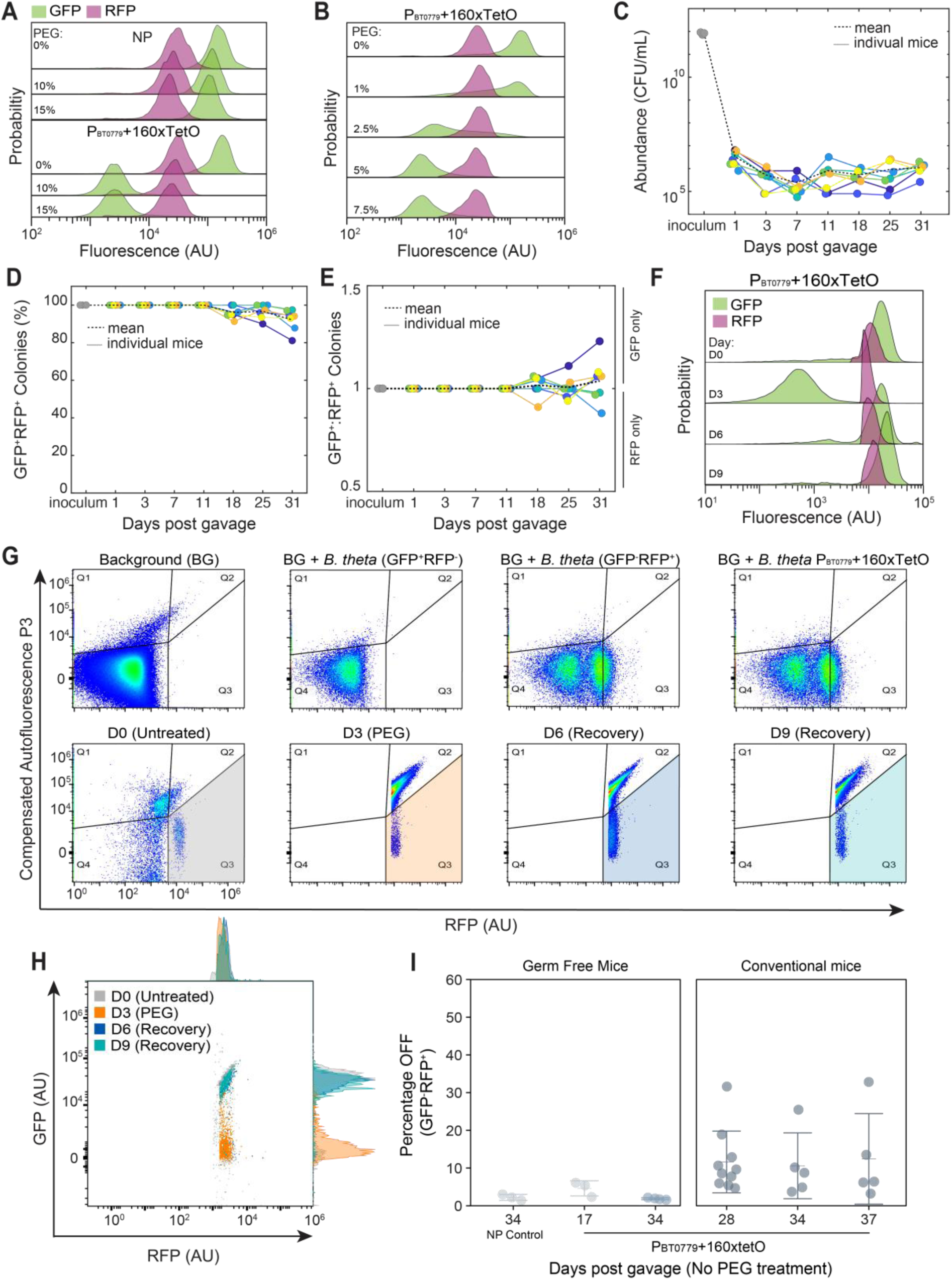
**Flow cytometry analysis of P_BT0779_+160xTetO biosensor fluorescence during PEG-induced malabsorption.** A) GFP and RFP fluorescence from cecal samples of mice colonized with NP or P_BT0779_+160xTetO biosensors after treatment with 0%, 10%, or 15% PEG (data from Figure 5A). B) GFP and RFP fluorescence from fecal samples of mice colonized with the P_BT0779_+160xTetO biosensor and treated with either 0%, 1%, 2.5%, 5%, or 7.5% PEG (data from Figure 5D). C) Colonization levels (in CFU/mL feces) of the P_BT0779_+160xTetO biosensor in individual mice hosting a complex microbiota over time. Colonies containing at least GFP or RFP fluorescence were counted from feces spot-plated on selective BHIS+Erm+Gent plates. Non-fluorescent colonies were identified as a different strain of *B. theta* by 16S rRNA sequencing. D) Maintenance of GFP^+^RFP^+^ reporters over time, from spot plating performed in C. E) Ratio of GFP^+^:RFP^+^ colonies over time, from spot plating performed in C. Ratios >1 indicate colonies that lost RFP, while ratios <1 indicate colonies that lost GFP. F) Background-subtracted GFP and RFP fluorescence from fecal samples of conventional mice colonized with the P_BT0779_+160xTetO biosensor collected prior to PEG treatment (D0), 3 days after 5% PEG treatment (D3), and 3- and 6-days post-PEG treatment (D6 and D9, respectively) as shown in Figure 5K. G) Flow cytometry gating strategy to identify biosensors from fecal samples of mice with a complex microbiota. *Top*: (*left to right*) Fecal samples from conventional mice (background; BG) were spiked with GFP^+^RFP^-^, GFP^-^RFP^+^, or an uninduced P_BT0779_+160xTetO biosensor strain grown *in vitro*. Gating was defined based on where RFP^+^ *B. theta* strains separated from BG autofluorescence in the P3 autofluorescence channel (Methods). *Bottom* (*left to right*): Gating was applied to samples from each day of PEG treatment Figure 5K; representative samples are shown here. H) GFP fluorescence from gated events in G (Q3; shaded) indicates biosensor activity only during PEG treatment. I) Proportion of biosensors lacking GFP fluorescence during PEG-free periods from experiments in (*left*) mono-colonization experiments in Figure 5A,D, and (*right*) the complex community experiment in Figure 5K. C-E: Solid lines and points indicate mean values from ∼10-40 single colonies counted from 7 individual mice; dashed lines indicate mean values for all colonies across all mice.

